# CoREST Complex Inhibition Alters RNA Splicing to Promote Neoantigen Expression and Enhance Tumor Immunity

**DOI:** 10.1101/2024.12.12.627852

**Authors:** Robert J. Fisher, Kihyun Park, Kwangwoon Lee, Katarina Pinjusic, Allison Vanasse, Christina S. Ennis, Scott Ficcaro, Jarrod Marto, Stephanie Stransky, Joseph Duke-Cohan, Anupa Geethadevi, Eric Raabe, Simone Sidoli, Chad W. Hicks, Derin B. Keskin, Catherine J. Wu, Philip A. Cole, Rhoda M. Alani

**Affiliations:** Department of Dermatology, Boston University Chobanian and Avedisian School of Medicine; Boston, Massachusetts, USA; Division of Genetics, Departments of Medicine and Biological Chemistry and Molecular Pharmacology, Harvard Medical School and Brigham and Women’s Hospital; Boston, Massachusetts, USA; Department of Medical Oncology, Dana-Farber Cancer Institute, Boston, Massachusetts, USA; Harvard Medical School; Boston, Massachusetts, USA; Cancer Center, Boston University Chobanian and Avedisian School of Medicine, Boston, Massachusetts, USA; Department of Cancer Biology, Center for Emergent Drug Targets, and Blais Proteomics Center, Dana-Farber Cancer Institute, Boston, MA, USA. Department of Pathology, Brigham and Women’s Hospital and Harvard Medical School; Department of Biochemistry, Albert Einstein College of Medicine; The Bronx, NY, USA; Division of Pediatric Oncology, Johns Hopkins University School of Medicine and Johns Hopkins Hospital, Bloomberg Children’s Center; Baltimore, Maryland, USA; Department of Pharmacology, Physiology & Biophysics, Boston University Chobanian and Avedisian School of Medicine; Boston, Massachusetts, USA; Broad Institute of MIT and Harvard, Cambridge, Massachusetts, USA; Translational Immunogenomics Laboratory, Dana-Farber Cancer Institute, Boston, Massachusetts, USA; Department of Computer Science, Metropolitan College, Boston University, Boston, Massachusetts, USA; Section for Bioinformatics, Department of Health Technology, Technical University of Denmark, Lyngby, Denmark

**Keywords:** CoREST, Melanoma, RNA splicing, Cryo-EM, Immunotherapy, Neoantigens

## Abstract

Epigenetic complexes tightly regulate gene expression and colocalize with RNA splicing machinery; however, the consequences of these interactions are uncertain. Here, we identify unique interactions of the CoREST repressor complex with RNA splicing factors and their functional consequences in tumorigenesis. Using mass spectrometry, in vivo binding assays, and cryo-EM we find that CoREST complex-splicing factor interactions are direct and perturbed by the CoREST complex inhibitor, corin, leading to extensive changes in RNA splicing in melanoma and other malignancies. Using predictive machine learning models and MHC IP-MS, we identify thousands of corin-induced neopeptides derived from unannotated splice sites which generate immunogenic splice-neoantigens. Furthermore, corin reactivates the response to immune checkpoint blockade and promotes dramatic expansion of cytotoxic T cells in an immune cold melanoma model. CoREST complex inhibition thus represents a unique therapeutic opportunity in cancer which creates tumor-associated neoantigens that enhance the immunogenicity of current therapeutics.

**Statement of Significance:** We identify a novel role of the CoREST transcriptional repressor complex in regulating pre-mRNA splicing and find that the small molecule inhibitor, corin, promotes alternative splicing events in cancer leading to neoantigen expression and T cell-mediated immunity. This represents a potential approach to promote immunoreactive neoantigen expression in immune-cold tumors.

## INTRODUCTION

Pre-mRNA splicing is essential for the expression of >95% of human genes which encode a diverse array of highly lineage and context-dependent protein variants^1,2^. Splicing changes are commonly seen in cancer, where tumors have been noted to have up to 30% more alternative splicing events (ASEs) than corresponding normal tissues, suggesting particular growth advantages of such changes^3,4^. The prevalence and impact of tumor-associated splicing changes have been recognized as a critical hallmark of cancer^5–7^ and splicing mechanisms have been implicated in resistance to cancer therapies^8,9^. Additionally, alternative RNA splicing increases proteomic diversity in tumors and induces expression of potential neoantigens for tumor-specific detection. These splice-neopeptides have been leveraged as immunotherapy targets^3,10–12^ and shown to elicit antigen-specific T cell activity to engage an anti-tumor response^13,14^; therefore, chemical induction of RNA splice modifications^15,16^ and therapies targeting pre-mRNA splicing in cancer are of great interest^11,15,17^.

Epigenetic macromolecular enzyme complexes tightly control gene expression at the chromatin level and have been shown to regulate transcript diversity through direct regulation of RNA splicing^18–25^. RNA splicing occurs largely cotranscriptionally, and alternative splice site choice is influenced by RNA polymerase II (Pol II) elongation rate, chromatin remodelers, and histone deacetylases^2,22,26^; however, the precise role of epigenetic complexes in RNA splicing is uncertain. During active transcription, histone modifying enzymes, nucleosome remodelers, general transcription factors, and the splicing machinery all localize within the same chromatin environment allowing for co-transcriptional crosstalk^20^ and epigenetic proteins have been found to modulate splicing by altering splicing factor gene expression^27,28^ and function^29^, interacting with spliceosomal and ribonucleoprotein complexes^30^, actively controlling the acetylation states of splicing-associated histone marks and splicing factors^30^, and altering novel splice junctions with transposable elements^31^.

The CoREST epigenetic repressor complex has core subunits HDAC1 (or close paralog HDAC2), LSD1 demethylase, and the scaffold protein CoREST and was originally identified as a corepressor complex for the transcription factor REST^32^. CoREST functions as a gene silencing complex through its histone deacetylase roles and demethylation of H3K4me^33,34^ and has been demonstrated to deacetylate the C-terminal domain (CTD) of the catalytic subunit of RNA Pol II as part of its transcriptional repressing effects^35^. We have previously described the dual LSD1/HDAC1 CoREST-selective inhibitor, corin^36^, and demonstrated its antineoplastic activity in melanoma^36,37^, diffuse glioma^38^, malignant peripheral nerve sheath tumor^39^, and colon cancer^40^. Here, we explore the role of the CoREST complex in RNA splicing regulation in melanoma. We uncover novel interactions between the CoREST complex and splicing factors, characterize these interactions using cryogenic electron-microscopy (cryo-EM) and define a noncanonical role for CoREST in RNA splicing regulation. We find that corin widely disrupts the CoREST-mediated splicing program in melanoma leading to induction of neoantigen expression in cell lines which is immunogenic. We further demonstrate that CoREST complex inhibition significantly reactivates the response to checkpoint blockade immunotherapy in immune cold tumors. We therefore suggest that CoREST complex inhibition represents a unique therapeutic opportunity in cancer.

## RESULTS

### The CoREST complex interacts with RNA splicing factors

The CoREST complex influences gene expression and is recruited to target genes through interactions with lineage-specific transcription factors, core transcription complex components and chromatin-associated proteins^33,41^ and has been shown by us and others to promote tumor growth in melanoma and other cancers^36–40^. In order to further define the mechanism of CoREST complex effects in melanoma, we evaluated its protein interactions in the setting of the CoREST inhibitor, corin (2.5 μM, 24h), versus DMSO control in two melanoma cell lines (1205Lu and 451Lu) (**Fig. 1A-D**; **Supplementary Table 1 and 2**). The LSD1/RCOR1 endogenous protein interactome was evaluated by liquid chromatography with tandem mass spectrometry (LC– MS/MS). LSD1 and RCOR1 were found to interact with all known CoREST complex subunits as well as the chromatin structural organizer, CTCF^24^, and several members of the SWI/SNF (BAF) complex, as has been previously described^42^. Interestingly, the CoREST complex-BAF interactions were disrupted by corin in both cell lines (**Supplementary Table 1**). Pathway analysis of proteins that had significant enrichment for LSD1 and RCOR1 binding over background across both cell lines notably identified significant enrichment for RNA splicing-related pathways, suggesting novel CoREST complex-splicing factor interactions in melanoma which are disrupted by corin (**Fig. 1B**).

**Fig. 1.**
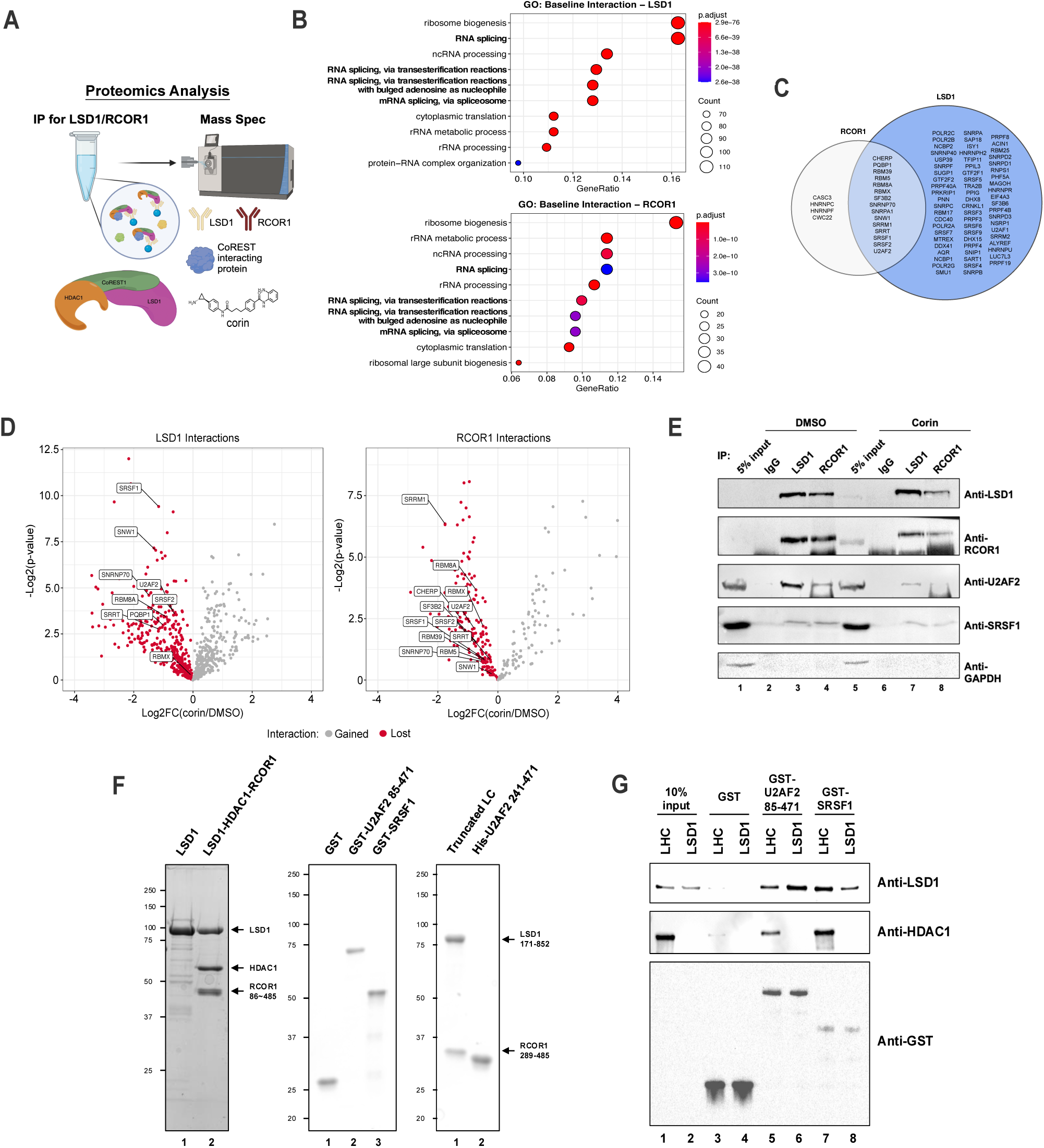
The CoREST complex interacts with splicing factors. **(A)** Overview of immunoprecipitation-mass spectrometry experiment completed in duplicate. **(B)** Gene ontology pathway analysis for proteins found to have significant baseline interactions (LogFC > 1, p < 0.05) with LSD1 (top) and RCOR1 (bottom). Enrichment analysis was performed using the hypergeometric test with multiple test correction by the Benjamini-Hochberg method. **(C)** Venn diagram of RNA splicing proteins found to significantly interact with LSD1 and RCOR1. **(D)** Volcano plots of LSD1 (left) and RCOR1 (right) baseline interactions lost with corin treatment (red). Labelled points are proteins from the overlap in (C). Statistical anlaysis was performed using the heteroscedastic t-test. **(E)** IP-WB analysis of CoREST complex-U2AF2 and CoREST complex-SRSF1 interactions with DMSO or corin treatment (24h, 2.5µM). **(F)** SDS-PAGE and Coomassie Blue staining of purified proteins. **(G)** GST pull-down assay using purified GST-tagged U2AF2 (amino acids 85–471) or SRSF1, and purified CoREST complex (LHC) or LSD1 protein.

To identify high confidence CoREST complex-interacting splicing factors (SFs), we overlapped SFs found in both LSD1 and RCOR1 pulldowns (**Fig. 1C**) then stratified the data by proteins that gain or lose interaction with the CoREST complex under corin treatment (**Fig 1D and Supplementary Table 2**). We identified 15 SFs that interacted with the CoREST complex (**Fig. 1C**) and found that corin disrupts 14 out of the 15 CoREST complex-SF interactions (**Fig. 1D**). To validate these findings, we selected two candidate SFs, U2 small nuclear RNA auxiliary factor 2 (U2AF2) and Serine/Arginine Splicing Factor 1 (SRSF1) based on the list of 15 CoREST complex-interacting SFs cross-referenced with an HDAC1 IP-MS dataset and their known functions in cancer,^43,44^ and performed IP-WB in an additional melanoma cell line (SKMEL5) (**Fig. 1E**). The CoREST complex was found to interact with both U2AF2 and SRSF1 by IP-WB and corin treatment greatly reduced CoREST-U2AF2 interactions (**Fig. 1E**). To determine whether the interaction between CoREST and U2AF2 is direct, we performed a GST pull-down assay using purified proteins (**Fig. 1F and G**). Full-length LSD1 protein was purified from a bacterial expression system, as well as the CoREST complex (LHC), which included LSD1, HDAC1, and RCOR1 (amino acids 86-485), from human 293F cells, and GST-tagged U2AF2 (amino acids 83-482) and SRSF1 from bacterial expression vectors (**Fig. 1F**). GST pull-down assays show that both LHC and LSD1 bind to U2AF2 and SRSF1 revealing direct interactions of LHC with U2AF2 and SRSF1, with LSD1 playing a key role in this interaction (**Fig. 1G**).

### Cryo-EM structure of the CoREST complex bound to U2AF2

In order to clarify the nature of CoREST complex interactions with the splicing machinery, we sought to develop structural details relevant to these complexes using cryo-EM analysis. Our studies were focused on the CoREST complex-U2AF2 interaction as U2AF2 displayed tighter binding with the CoREST complex in AlphaFold^45^ predictions compared to SRSF1. We determined the cryo-EM structure of U2AF2 bound to LSD1 + RCOR1 **(Supplementary Fig. 1A)** at a global resolution of 5.14 Å **(Supplementary Fig. 1B)**.

To prepare cryo-EM samples, we purified the LSD1 (a.a. 171-852) and RCOR1 (a.a. 286-485) complex (truncated LC) and U2AF2 (a.a. 241-471) proteins separately (**Fig. 1F**), mixed truncated LC with U2AF2 (a.a. 241-475) at a 1:5 ratio in the absence of crosslinker and performed size-exclusion purification to isolate the U2AF2-bound LSD1/RCOR1 complex (**Fig. 2A**). The chromatogram showed a peak corresponding to the LSD1-RCOR1-U2AF2 in complex, as well as two separate peaks for LSD1-RCOR1 and U2AF2 alone, confirming that the LSD1-RCOR1 complex binds U2AF2. Fractions containing the LSD1-RCOR1-U2AF2 complex were pooled (22-25) and samples were frozen onto cryo-EM grids. Of note, the U2AF2 (a.a. 241-475) construct used for the cryo-EM sample contains two structured globular domains, the RNA Recognition Motif 2 (RRM2) and U2AF2 Homology Motif (UHM) (**Fig. 2B**).

**Fig. 2.**
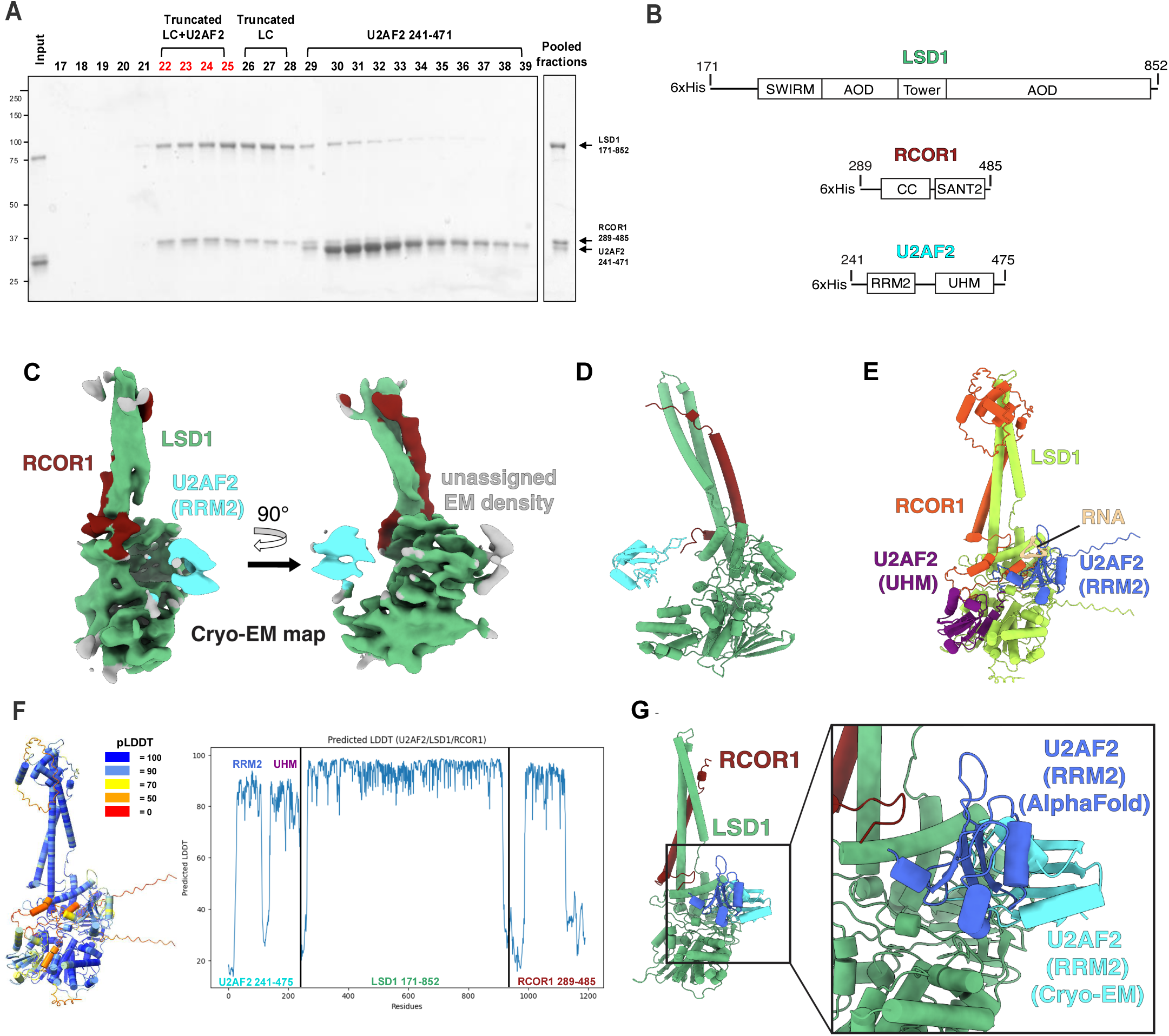
Cryo-EM structure and AlphaFold prediction of U2AF2 bound to LSD1+RCOR1. **(A)** Size exclusion chromatography for cryo-EM sample preparation. The early fractions (fractions 22–25), containing LSD1, RCOR1, and U2AF2, were pooled, concentrated and subsequently used for cryo-EM analysis. **(B)** Domain schematic of all protein components used for cryo-EM sample preparation. **(C)** Cryo-EM map of the RRM2 domain of U2AF2 bound to the LSD1+RCOR1 complex. **(D)** Cryo-EM model in cartoon view representing the cryo-EM map shown in panel (C). (**E**) AlphaFold model representing the prediction shown in panel (D). **(F)** Predicted local distance difference test (pLDDT) plot of the AlphaFold multimer prediction incorporating LSD1, RCOR1, U2AF2, and RNA. **(G)** Superimposition of the RRM2 domain of U2AF2 from AlphaFold multimer over our cryo-EM structure of LSD1+RCOR1+U2AF2. Zoomed-in box shows the degree of similarity in position between the two RRM2 globular domain models.

We observed well-resolved EM density for the LSD1+RCOR1 complex, low-resolution density for U2AF2 adjacent to the side of LSD1, and some unassigned very-low resolution density in contact with the opposite side of LSD1 (**Fig. 2C**). The resolution of the EM density corresponding to LSD1+RCOR1, at ∼5.0 – 6.0 Å **(Supplementary Fig. 1C)**, was sufficient to unambiguously rigid fit the crystal structure of the LSD1+RCOR1 complex (PDB: 2IW5)^46^ within the EM map **(Supplementary Fig. 1D)**. Therefore, our cryo-EM structure, the first cryo-EM structure of the LSD1+RCOR1 complex, appears to adopt a similar conformation to the crystallized form of the LSD1+RCOR1 complex^46,47^.

The resolution of the adjacent U2AF2 EM density, at 6.0 – 7.0 Å, was found to be lower than the LSD1+RCOR1 complex **(Supplementary Fig. 1C)**. We therefore compared the relative size of the two U2AF2 globular domains within the EM map **(Supplementary Fig. 1E and F)** and determined that the adjacent U2AF2 density is likely to be the RRM2 domain of U2AF2, as opposed to the UHM domain. The resolution for the RRM2 EM density was not high enough to determine an accurate orientation of the model within the EM map, likely due to its extensive conformational heterogeneity; however, we were able to determine that the structured RRM2 domain does not directly contact LSD1 (**Fig. 2C and 2D**). The additional EM density contacting LSD1 opposite the RRM2 domain could not be assigned (**Fig. 2C**), although it is possible that the unassigned density may represent the UHM domain of U2AF2.

To better understand the mechanism of RNA recognition by U2AF2 in the context of the LSD1+RCOR1+U2AF2 complex, we performed an AlphaFold3^45^ multimer calculation of LSD1+RCOR1+U2AF2+RNA. As an input, we provided the same sequences as the cryoEM constructs LSD1, RCOR1, and U2AF2 plus added an additional 11 base-pairs of RNA **(see Methods)**. The AlphaFold prediction revealed a structured LSD1+RCOR1 at very-high confidence and showed the RRM2 and UHM domains of U2AF2 bound to LSD1 at moderately high confidence (**Fig. 2E and 2F**). The predicted local distance difference test (pLDDT), representing local structural confidence calculated by AlphaFold, shows that the two globular domains of U2AF2 (RRM2 and UHM) have high scores, while the flexible linker loop connecting them has low scores (**Fig. 2F**) demonstrating AlphaFold prediction of U2AF2 remaining folded in globular domains and associated with LSD1+RCOR1 as observed in the cryo-EM structure. Superimposing the AlphaFold structure of the RRM2 domain of U2AF2 over our cryo-EM structure shows the AlphaFold model of RRM2 in close proximity to the cryo-EM model of RRM2 (**Fig. 2G**). The AlphaFold prediction also revealed RNA bound to the RRM2 domain of U2AF2 (**Fig. 2E**), a binding mode that has been previously observed in crystal structures of RNA bound to U2AF2^48,49^. While we did not include RNA in our cryo-EM sample, the position of the RRM2 domain of U2AF2 could also accommodate bound RNA.

### CoREST complex regulation of splicing factor gene expression

Given our finding of the CoREST complex binding to components of the RNA splicing machinery which is lost in the context of the CoREST inhibitor, corin, and the fact that expression of RNA binding proteins (RBPs) is widely dysregulated in solid tumors^50^, we sought to determine whether CoREST may directly impact the transcription of pre-mRNA splicing factors. We conducted RNA-sequencing on six melanoma cell lines treated with corin (2.5μM, 24h), three tumor cell lines designated as MITF^high^/AXL^high^ cell lines (differentiated phenotype)^51^ and three tumor cell lines designated as MITF^low^/AXL^high^ (dedifferentiated phenotype)^51^, and performed gene set enrichment analysis (GSEA) to identify commonly enriched pathways following corin treatment across all cell lines regardless of phenotypic status. Remarkably, we identified the KEGG Spliceosome pathway as the most common downregulated pathway across all six cell lines (**Fig. 3A and 3B**) suggesting that CoREST inhibition downregulates splicing factor gene expression in melanoma regardless of molecular phenotype. Further analysis of the splicing factor genes impacted by corin treatment revealed that, although many U2-related and early spliceosomal genes were downregulated, a broad range of spliceosome and trans-acting splicing regulators were also downregulated suggesting widespread impact on the splicing machinery (**Fig. 3C and Supplementary Fig. 2A**). Notably, 29% of genes in the KEGG Spliceosome gene set were found to be consistently downregulated by corin without correlation to melanoma phenotype highlighting the broad impact of CoREST on splicing factor gene regulation. To validate these findings, western blotting was performed on the three most downregulated genes where U2AF2 and ALYREF were both noted to be significantly decreased in expression at the protein level following corin treatment (**Fig. 3D, 3E, and Supplementary Fig. 2B**). This is particularly of interest since melanoma (TCGA SKCM) patients with decreased U2AF2 expression have significantly prolonged survival (p=0.001; **Fig. 3F**).

**Fig. 3.**
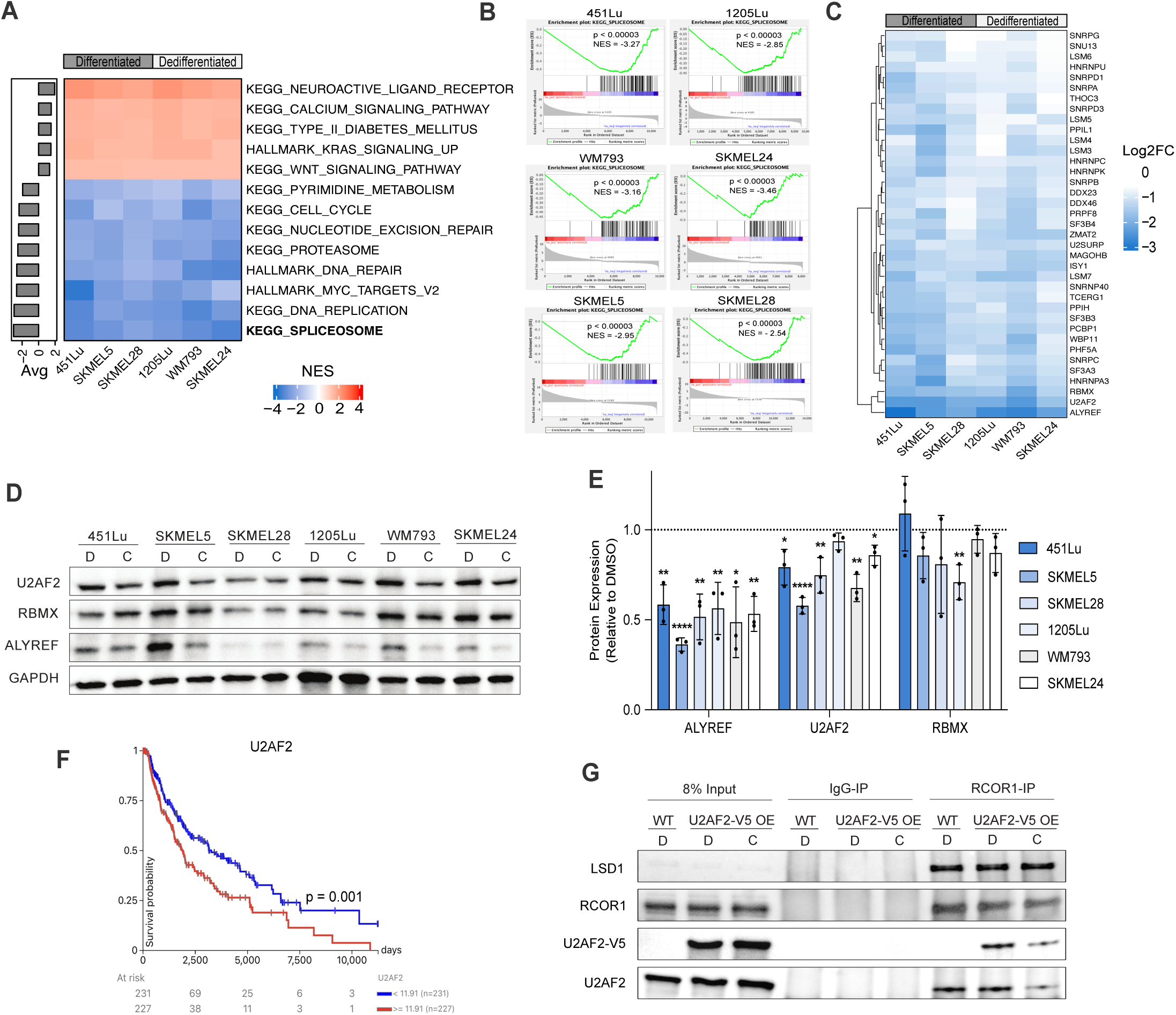
CoREST transcriptionally regulates splicing factor gene expression. **(A)** Heatmap of significant KEGG and Hallmark pathways (nominal p < 0.05) across six melanoma cell lines treated with corin (24h, 2.5µM) in duplicate. Pathways are ranked by average normalized enrichment score (NES) and cell lines are grouped based on phenotype. **(B)** Gene Set Enrichment Analysis plots for each cell line showing a significant negative enrichment for "KEGG Spliceosome". **(C)** Heatmap of splicing factor genes significantly downregulated by corin treatment (q < 0.01, Log2FC < -0.5) across all six melanoma cell lines clustered using Euclidean distance. **(D)** Representative western blot of downregulated splicing factors across six melanoma cell lines treated with DMSO (D) or corin (C) and quantification of biological replicates (n =3) **(E).** Error bars represent the standard deviation (SD). **(F)** Kaplan-Meier plot of TCGA-SKCM patient survival based on median U2AF2 expression. Significance was determined using a log-rank test. **(g)** IP-WB analysis of CoREST-U2AF2 interactions following DMSO (D) and corin (C) treatment (24h, 2.5µM) in V5-tagged U2AF2 overexpression SKMEL5 cells.

To determine whether corin-associated decreases in splicing factor expression were due to direct transcriptional changes, we performed PRO-seq on SKMEL5 cells treated with corin and used the 3’ nascent RNA reads to conduct differential gene expression analysis (**Supplementary Fig. 2C**). We found the KEGG Spliceosome pathway to be significantly negatively enriched in the setting of corin treatment with over 50% of splicing factor genes downregulated in the RNA-seq also downregulated in the PRO-seq (**Supplementary Fig. 2D**). This suggests that corin-associated splicing factor downregulation is due to direct effects on mRNA synthesis rather than transcript stability. As CoREST is an epigenetic repressor complex and the vast majority of corin effects on tumor cells are associated with transcriptional upregulation^36–39^, it is anticipated that these transcriptional repressive effects of corin on splicing factor genes are associated with indirect transcriptional changes. Since we were able to demonstrate decreased transcription of SF genes in the setting of corin in melanoma cells, we sought to determine whether the loss of CoREST complex-SF protein interactions in the setting of corin was due to decreased expression of splicing factor proteins, or a disruption of CoREST complex-SF binding. Using V5-tagged U2AF2 overexpression in SKMEL5 melanoma cells, we found that binding of V5-U2AF2 to CoREST (RCOR1) was significantly inhibited in the setting of corin treatment without a change in V5-U2AF2 expression supporting a specific inhibitory effect of corin on U2AF2 binding to CoREST independent of the transcriptional effects on U2AF2 (**Fig. 3G**).

### Corin induces RNA splicing changes in melanoma

Given the observed CoREST complex-U2AF2 interactions and corin’s impact on splicing factor gene expression, we next sought to determine the functional effects of corin on RNA splicing across a panel of melanoma cell lines. Two computational tools were used to analyze our RNA-seq data for changes in splicing events. rMATS^52^ was applied to detect alterations in skipped exons (SE), 3’ splice sites (3’SS), 5’ splice sites (5’SS), and mutually exclusive exons (MXE), while LeafCutter^53^ was used to identify changes in retained introns (RI). Using these methods, we identified thousands of splicing events modified by corin treatment across the six melanoma cell lines tested (**Fig. 4A and Supplementary Table 3**), with skipped exons being the most frequently affected. When we compared PSI (percent spliced in) values of SE events between DMSO and corin treatment, all cell lines except WM793 showed a significant decrease in PSI distribution in the setting of corin treatment indicating increased exon skipping with CoREST complex inhibition (**Fig. 4B**). Additionally, when we compared the exon architecture of skipped versus included exons, skipped exons were significantly shorter, flanked by long introns, and in lower GC content regions of the genome (**Supplementary Fig. 3A and B**). This suggests that the CoREST complex may be involved in promoting exon inclusion preferentially at defined exon architecture during co-transcriptional RNA splicing.

**Fig. 4.**
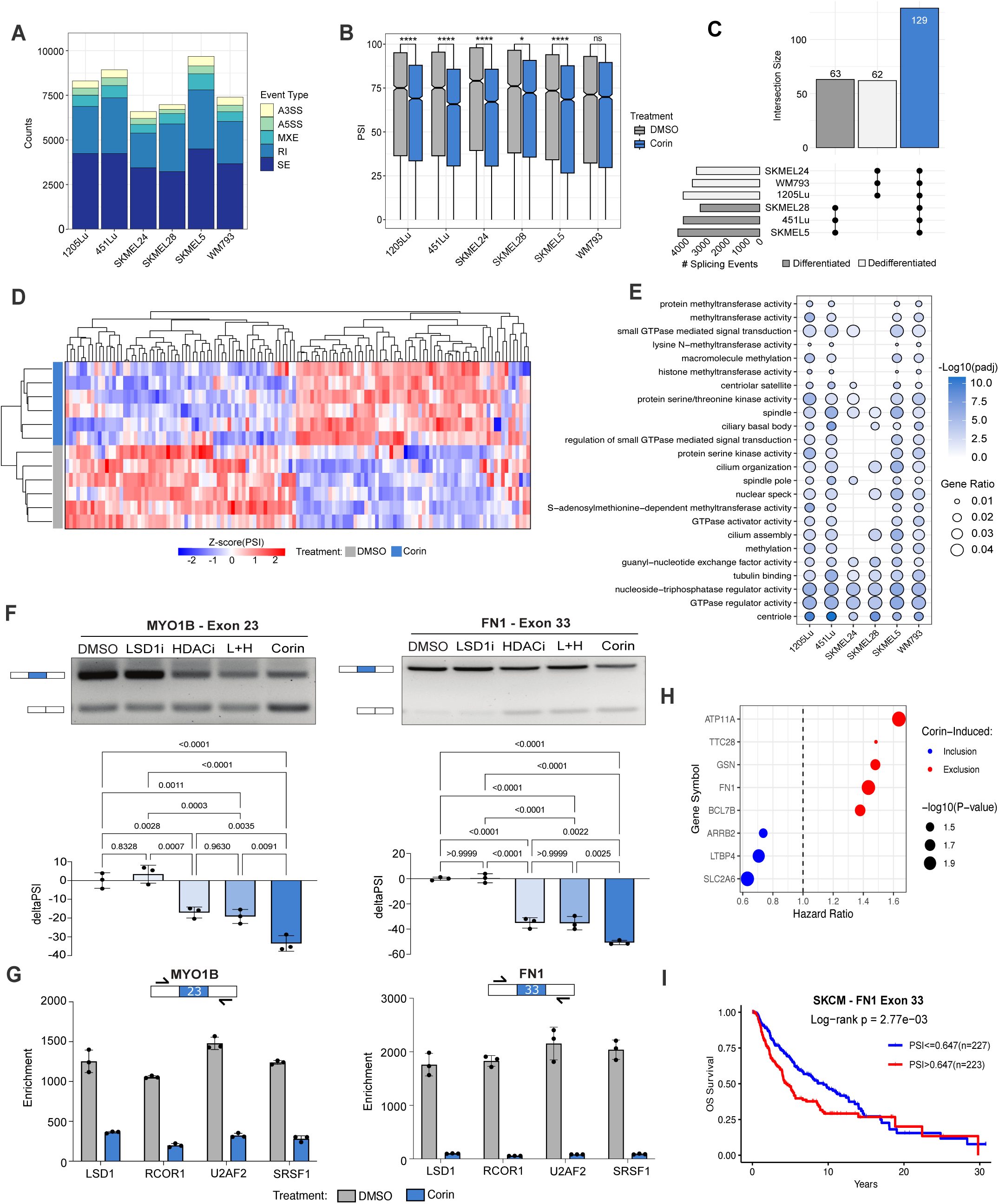
Corin induces RNA splicing changes in melanoma. **(A)** Summary of significant RNA splicing changes across six melanoma cell lines treated with corin (deltaPSI ≥ |0.1|, q < 0.05) in duplicate. **(B)** Percent Spliced In (PSI) levels for all significant SE events following DMSO and corin treatment. Statistical comparisons were performed using a two-sample t-test to assess differences in PSI value between treatment groups within each cell line. P-values were adjusted for multiple comparisons using the Bonferroni correction (*p.adj < 0.05, **p.adj < 0.01, ***p.adj < 0.001, ****p.adj < 0.0001). **(C)** UpSet plot of skipped exon (SE) events that are exclusive to the differentiated phenotype, dedifferentiated phenotype, or shared by all cell lines (blue). **(D)** Unsupervised hierarchical clustering heatmap based on Euclidian distance of shared skipped exon inclusion levels. Rows are melanoma cell lines clustered by treatment and columns are shared inclusion events. **(E)** Gene ontology dotplot of the top pathways impacted by corin-induced differential exon inclusion across all cell lines (median padj < 0.01). Enrichment analysis was performed using the hypergeometric test with multiple test correction by the Benjamini-Hochberg method. **(F)** Representative RT-PCR gels and quantification comparing corin to single agent inhibitors of HDAC and LSD1 in MYO1B and FN1 splicing. Statistical analysis of biological replicates (n=3) was performed using an unpaired, two-tailed t-test. Error bars represent the standard deviation (SD). **(G)** Representative RIP-qPCR biological replicate (n=2) with 3 technical replicates of CoREST complex subunits and splicing factor occupancy at MYO1B and FN1 splice sites with corin treatment. Error bars represent the standard deviation (SD). **(H)** Hazard ratios for corin-induced splicing events based on TCGA-SKCM survival data measured using the Cox proportional hazards regression. **(I)** Kaplan-Meier curve of FN1 exon 33 inclusion in TCGA-SKCM using the median PSI threshold. Significance was determined using a log-rank test.

Next, we compared the unique SE events stratified by MITF status to determine whether the CoREST complex may be responsible for driving a phenotypic splicing program (**Fig. 4C**). We identified several phenotype-specific SE events, with 63 events associated with the differentiated (MITF^high^) state and 62 associated with the dedifferentiated (MITF^low^) state; however, the overlap was minimal relative to the total number of SE events across the cell lines. We did, however, find 129 SE events that were commonly induced by corin treatment across all melanoma cell lines which may be more generalizable targets of CoREST complex-specific splicing (**Fig. 4D**). Although, these common SE events comprise a small fraction of the total, gene ontology analysis of corin-induced differential exon inclusion conducted independently on each cell line demonstrated that many of the same biological processes were impacted by corin treatment such as cell motility, cytoskeletal structure, and GTPase activity (**Fig. 4E**).

Given the impact of CoREST complex inhibition on SF binding to the CoREST complex and SF transcription, we next sought to determine the effect of corin on constitutive pre-mRNA splicing activity by comparing the PSI distribution of intron retention events identified by LeafCutter and found no significant directional increase in retention in constitutively spliced introns. We further compared the pathways impacted by intron retention to those impacted by exon skipping and found that RNA splicing factors are most differentially impacted by corin-induced intronic splicing (**Supplementary Fig. 3C and D**). In order to define the splicing factors that drive the CoREST complex’s impact on RNA splicing, we utilized RNA-SPRINT^54^ to estimate individual RNA Binding Protein (RBP) activity across all six cell lines under each treatment condition and correlated the activity scores to the expression level of the corresponding RBP using Spearman’s rank correlation (**Supplementary Fig. 4A**). Four candidates (SF3B4, HNRNPC, HNRNPK, and U2AF2) had significant coefficients (ρ > 0.5) in which decreased expression was strongly correlated with decreased RBP activity (**Supplementary Fig. 4B-D**), suggesting that CoREST complex-induced splicing programs could be largely attributed to the functions of these particular splicing factors. Overall, these findings highlight the complexity of the systems in place to critically regulate RNA splicing, as we see the CoREST complex engaged in transcriptional, translational, and post-translational modifications that control the splicing process as well as specific splicing of splicing-associated transcripts.

Next, we validated 3 common SE events associated with cytoskeletal structure and cell motility functions with demonstrated oncogenic associations by RT-PCR (**Fig. 4F**; **Supplementary Fig. 5A-C**). Myosin-1b (MYO1B) is a motor protein critical for actin filament organization and is crucial for neuronal development. MYO1B exon 23 exclusion has been associated with decreased cell migration and significant survival outcomes in glioblastoma mouse models^28^. Fibronectin (FN1) exon 33 encodes extra domain A which has been shown to increase tumor metastasis and is widely expressed in melanoma tissues compared to normal skin^55^. Tight Junction Protein 1 (TJP1 or ZO-1) exon 20 encodes the alpha domain whose skipped isoform is increased in breast, lung, and colon cancers^56^ and has been shown to enhance actin stress fiber assembly, increase cell migration, and is induced during EMT by TGFB^56^. In all instances, we found that corin significantly reversed tumor-associated splice isoforms (**Fig. 4F**; **Supplementary Fig. 5A-C**); moreover, we found that corin significantly reversed CoREST complex-mediated splicing events to a greater extent than either MS-275 or GSK-LSD1 alone, or the combination of inhibitors (**Fig. 4F**; **Supplementary Fig. 5A-C**) suggesting targeted impacts of the dual-specificity CoREST complex inhibitor.

Based on our finding that CoREST complex-U2AF2 interactions are disrupted in the setting of corin, we hypothesized that these differential exon inclusion events may be directly influenced by CoREST-SF recruitment to RNA. Using RIP-qPCR with primers flanking FN1 exon 33 and MYO1B exon 23, we found that CoREST complex subunits (LSD1 and RCOR1) and SFs (U2AF2 and SRSF1) not only bind RNA at these splice sites but also significantly lose occupancy with corin treatment (**Fig. 4G**), suggesting that CoREST complex-SF recruitment to alternative splice sites can directly impact exon inclusion levels.

Given the impact of corin on tumor-associated splicing events, we analyzed the 129 corin-induced common SE events in the 6 melanoma cell lines for survival-association in TCGA-SKCM (human skin cutaneous melanoma) using OncoSplicing and a cox regression model^57^. Eight of the common splicing events were found to be directionally beneficial for overall survival across TCGA SKCM patients (**Fig. 4H**) including FN1 exon 33, where exon exclusion was found to be associated with significantly improved survival outcomes (**Fig. 4I**).

As mRNA synthesis rates have been shown to have a significant impact on splicing, we were curious to see if CoREST could regulate RNA splicing by influencing the rate of active transcription. A previous study found that the CoREST complex can directly bind to RNA polymerase II after the pre-initiation complex is formed to deacetylate the CTD and effectively pause the polymerase^35^. We hypothesized that this noncanonical role of the CoREST complex could alter RNA synthesis kinetics, and thus alter exon inclusion levels. To test this, we analyzed our PRO-seq data for changes in Pol II coverage at alternatively included exon splice sites in the setting of corin. Although we found that inhibiting the CoREST complex led to significant promoter-proximal pause release genome-wide (**Supplementary Fig. 6A and B**), we did not note a significant difference in Pol II coverage between included or excluded exons (**Supplementary Fig. 6C**), suggesting that these alternative splicing changes occur independent of a corin-induced kinetic effect on Pol II pause release.

### Corin induces RNA splicing changes across cancers

Given the broad impact of corin on the splicing machinery in melanoma, we were interested in determining whether corin could modulate splicing across other cancers. Comparison of the expression levels of splicing factors transcriptionally downregulated by corin treatment between normal and matched tumor samples in cBioPortal revealed that many of these factors are significantly overexpressed in a diverse array of tumors (**Fig. 5A**), suggesting that this may be a driver event in cancers which could be targeted by corin. We next sought to determine whether corin-induced splicing changes could be seen in other cancers. Data obtained from two independent external RNA-sequencing datasets using cancer cells treated with corin, an ER+ breast cancer dataset^42^ and an Atypical Teratoid Rhabdoid Tumor (ATRT) dataset (unpublished results), were mined for gene expression changes and differential splicing analysis was performed using the pipeline previously described. Remarkably, we found that corin significantly impacted splicing factor gene expression within these tumor cells to a similar extent to that seen in melanoma (p < 0.001; **Fig. 5B and C**) and promoted thousands of RNA splicing differences across both tumor types (**Fig. 5D and Supplementary Table 4**). Moreover, the pathways affected by corin-induced splicing included many of the same cell motility and GTPase pathways seen in melanoma (**Fig. 5E and F**) suggesting not only that corin can induce splicing changes across cancer types, but also that the CoREST complex may be involved in regulating splicing of specific biological processes in cancer.

**Fig. 5.**
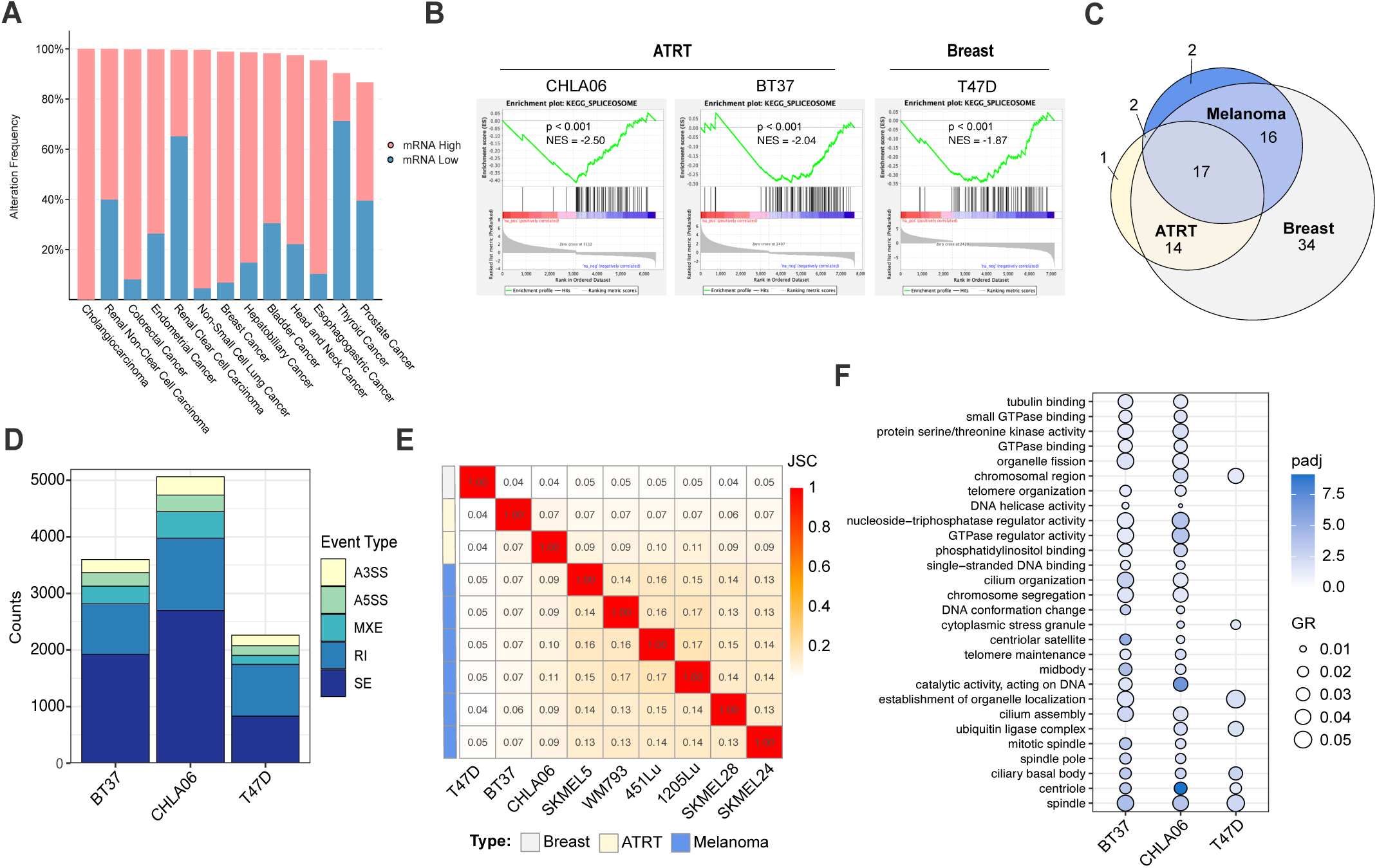
Corin impacts RNA splicing across cancers. **(A)** Histogram depicting the frequency of altered mRNA expression alteration in CoREST complex-regulated RNA splicing factors between cancer and matched normal tissue from cBioPortal stratified by cancer type. Pink bars indicate the frequency of overexpression events and blue bars indicate the frequency of downregulation events. **(B)** Gene Set Enrichment Analysis plots for “KEGG Spliceosome” in ATRT and breast cancer cell lines treated with corin. **(C)** Venn diagram of splicing factors significantly downregulated by corin in melanoma, ATRT, and breast cancer. **(D)** Summary of corin-induced RNA splicing changes in ATRT and breast cancer cells. **(E)** Jaccard similarity index comparing corin-induced splicing events across all cell lines and cancer types. **(F)** Gene ontology analysis of common significant pathways affected by corin-induced SE splicing events in ATRT and breast cancer. Enrichment analysis was performed using the hypergeometric test with multiple test correction by the Benjamini-Hochberg method.

### Corin-induced splice-neoantigens are presented on human MHC and are immunogenic

The splicing events discussed thus far have arisen from known annotations; however, we found that many events induced by corin were derived from unannotated splice sites and hypothesized that these unannotated events would have the potential to produce neopeptide products that may elicit an immune response in vivo^15,16^. To identify neopeptide candidates, we utilized a combination of predictive computational methods and HLA IP-mass spectrometry validation (**Fig. 6A**). SpliceTools^58^ and SNAF^54^ were used to identify 8-11mer neopeptides produced from significant splicing events (q < 0.05, |deltaPSI| ≥ 0.1, log2TPM > 3). Remarkably, we identified thousands of neopeptide sequences induced by corin treatment across all melanoma cell lines and found them to be induced in a cell-line dependent manner (**Fig. 6B and C**) akin to the cell-line specific splicing patterns identified previously (**Fig. 4**). In order to assess HLA binding of predicted neopeptides we utilized two machine learning models trained on patient mass spectrometry data: NetMHCpan4.1^59^ and HLAthena^60^. We found hundreds of neopeptides predicted to bind (%Rank < 2) SKMEL5 cell HLAs by both tools (**Fig. 6D**) and overlapped the neopeptides identified by each tool for each allele to identify the best candidate peptides (**Fig. 6E**). Corin-associated neopeptides were then ranked based on a scoring system taking HLA binding rank, junction count, and deltaPSI value into consideration (**Fig. 6F and Supplementary Table 5**). HLA-IP mass spectrometry was used to validate the 556 candidate neopeptide targets. Remarkably, we recovered over 10,000 8-11mer peptides in each replicate of mass spec and found 7462 corin-specific peptide products (**Fig. 6G and Supplementary Table 6**). When we overlapped those products with the predicted 556 candidate neopeptides, we found 2 neopeptides that were produced from corin-specific splicing (**Fig. 6H**). To assess if any neopeptides were missed, we also overlapped the 7462 corin-specific peptides with the human proteome and found the 2 neopeptides already predicted. Lastly, we used the deep learning model, DeepImmuno^61^, to predict the immunogenicity score of each peptide to its respective strongest binding HLA allele. Both peptides validated by mass spec were predicted to be immunogenic (immunogenicity score > 0.5) and were further tested in immunogenicity assays. Additionally, we ranked the top predicted peptides based on immunogenicity score and tested the top two unique peptides in our immunogenicity assay (**Fig. 6I**). Since HLA-C alleles do not have adequate training data for accurate DeepImmuno immunogenicity predictions, we focused on HLA-A and HLA-B binding peptides.

**Fig. 6.**
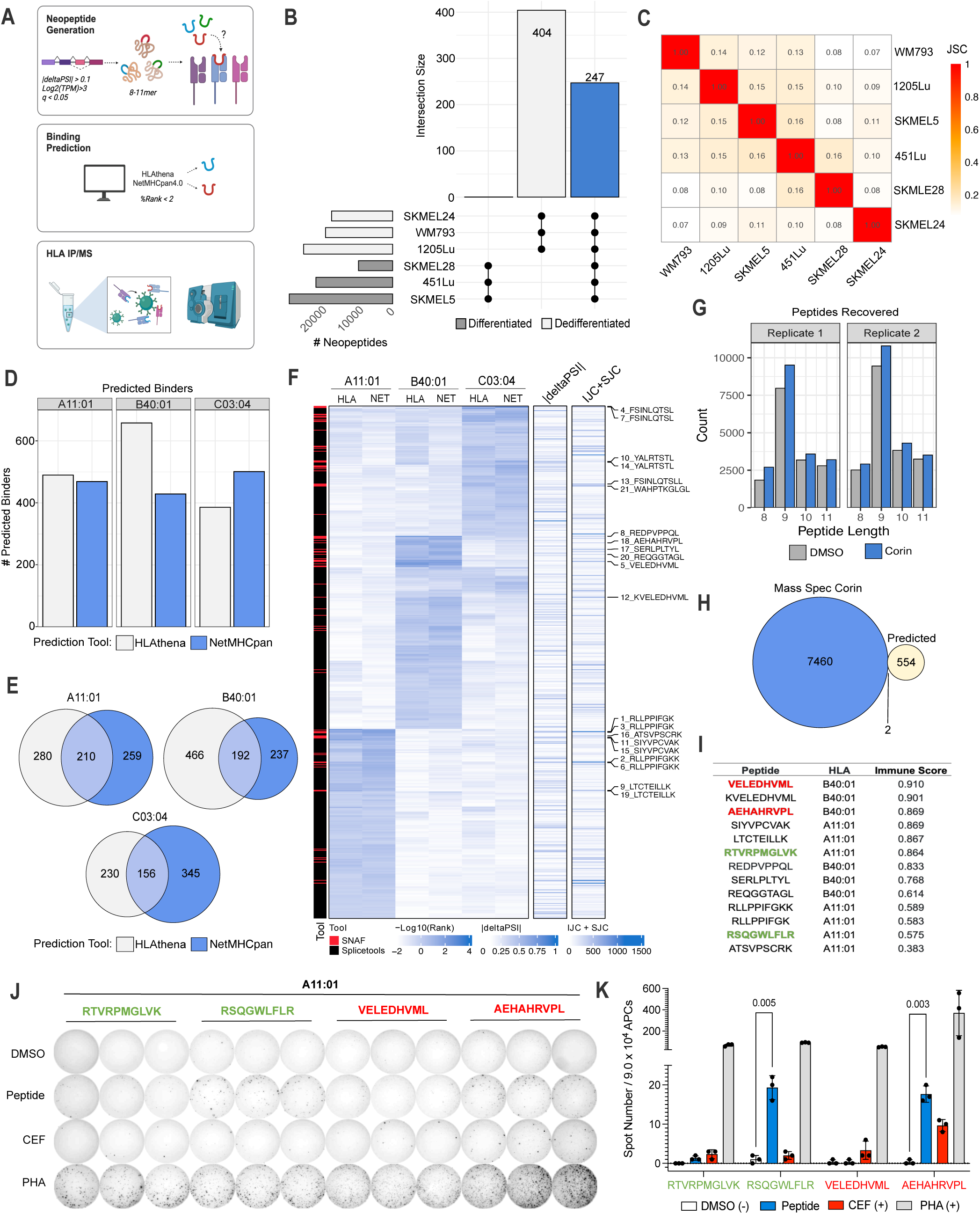
Corin-induced splicing produces neoantigens that bind human MHC and are immunogenic. **(A)** Overview of neopeptide discovery and MHC binding predictions **(B)** UpSet plot of neopeptides (8-11mers) produced with corin treatment of melanoma cells that are exclusive to the differentiated phenotype, dedifferentiated phenotype, or shared by all cell lines (blue). **(C)** Jaccard similarity index comparing corin-induced neopeptide production across all melanoma cell lines. **(D)** Number of corin-induced neopeptides predicted to bind to SKMEL5 HLAs based on two prediction tools: HLAthena and NetMHCPan4.1 (%Rank < 2). **(E)** Overlap of SKMEL5 corin-induced neopeptide HLA binders for each allele predicted by both tools. **(F)** Heatmap showing binding scores, PSI values, and junction counts of predicted SKMEL5 corin-induced neopeptides identified by SNAF and Splicetools. The top 15 unique candidates are labeled. **(G)** Histogram plots of peptides recovered from MHC-IP/MS in SKMEL5 DMSO and corin-treated (72h, 1µM) samples for each replicate (n=2). **(H)** Identification of SKMEL5 corin-induced neopeptides recovered by MHC IP-MS. Corin-exclusive peptides are those identified from the IP-MS that appear in at least one corin replicate but neither DMSO replicate. Predicted peptides are those identified by binding scores in (F). **(I)** Immunogenicity score predictions for neopeptide candidates. Green peptides are those identified from IP-MS and red peptides are additional candidates selected for immunogenicity validation assays based on immunogenic prediction and scores from (F). **(J)** Ex vivo IFNγ ELISpot assay for each candidate neopeptide tested with CEF and PHA positive controls. HLA-matched PBMCs were pre-stimulated with synthesized peptides (10µg/ml) for 14 days in IL2/IL7 media. APCs were isolated from CD4 and CD8 depleted PBMCs, loaded with peptides (10µg/ml), and seeded at a ratio of 3:1 with the pre-stimulated T cells in a 96 well ELISpot plate and analysed for IFNγ + T cells. **(K)** Quantification of ex vivo IFNγ ELISpot assay illustrated in (J). Statistical analysis was performed using multiple two-tailed unpaired t-tests. Error bars represent the standard deviation (SD).

To test the immunogenic potential of the four candidate neoantigens, we loaded high purity synthetic peptides on antigen presenting cells (APCs) derived from HLA-matched PBMCs (A11:01, B40:01, C03:04) and assessed pre-stimulated CD8+ T cell activation in an IFN-y Enzyme-Linked ImmunoSpot (ELISpot) assay (**Fig. 6J, K**; **Supplementary Fig. 7A and B**). Two of the four neopeptides elicited strong T cell activation when loaded on A11:01 APCs compared to a DMSO negative control and CEF pool positive control confirming that corin treatment can induce splice-neoantigens in human melanoma cells that are immunogenic.

### Corin treatment sensitizes immune cold tumors to immunotherapy

Given our finding of corin-induced expression of immunogenic neoantigens, we sought to evaluate corin treatment of melanoma in an immune competent mouse model in conjunction with immune checkpoint blockade (ICB) in an immune “cold” melanoma model. The B16-F10 melanoma mouse model was established with treatment arms including vehicle, ⍺-PD1, corin, and the combination of ⍺-PD1 + corin (**Fig. 7A**) and tumors were measured starting 7 days after inoculation. Remarkably, the combination ⍺-PD-1 + corin treatment was found to decrease tumor growth by 66% as measured by tumor volume and tumor weight compared to ⍺-PD1 treatment alone within one week of initiating therapy (**Fig. 7B, C, and D**). Additionally, there were no significant changes in body (**Fig. 7E**) or spleen weight (**Fig. 7F**) suggesting that corin treatment is tolerated at the treatment dose. In order to assess the impact of corin treatment on the immune microenvironment, we isolated CD45+ cells from ⍺-PD1 and ⍺-PD1+corin tumors and evaluated them by single-cell RNA sequencing (scRNA-seq) (**Fig. 7G-L**; **Supplementary Fig. 7C-F**). 23,000 tumor-enriched CD45+ cells were profiled, analyzed by Uniform manifold approximation and projection (UMAP), and segregated into 17 clusters based on the expression of leukocyte-associated genes and enrichment of gene signatures from external single-cell datasets (**Fig. 7G-J**)^62^. This process allowed us to define major canonical T cell phenotypes such as naive T cells (Tn), exhausted cells (Tex), CD8+ cytotoxic T lymphocytes (CTLs), regulatory T cells (Treg), and cycling cells (Tcyc) as well as monocyte/macrophage lineages, B cells, neutrophils, dendritic cells and plasma cells (**Supplementary Fig. 7C and D**). Strikingly, T cells from the ⍺PD-1 treated tumors were primarily Tn cells (43%) and Tex (19%) whereas T cells from the corin+ ⍺PD-1-treated tumors underwent significant expansion of CTLs (57%) which supports induction of a CTL response by tumor-associated neoantigens (**Fig. 7I, J**, **and Supplementary Fig. 7E**). Differential gene expression analysis in the T cell populations from ⍺PD-1 versus ⍺PD-1+corin treated tumors showed significant upregulation of granzyme B (Gzmb), interferon gamma (Ifng) and Cd8a along with significant downregulation of transcription factor 7 (Tcf7), lymphoid enhancer binding factor 1 (Lef1), and L-selectin (Sell) (**Fig. 7K and Supplementary Fig. 7F**). Gene set enrichment analysis (GSEA) demonstrated enhanced cytokine activity, leukocyte migration inflammatory response, antigen response, and tumor-associated immune response in T cells isolated from ⍺PD-1+corin treated tumors versus ⍺PD-1 treatment alone (**Fig. 7L**), consistent with T-cell activation in response to tumor-associated neoantigen expression.

**Fig. 7.**
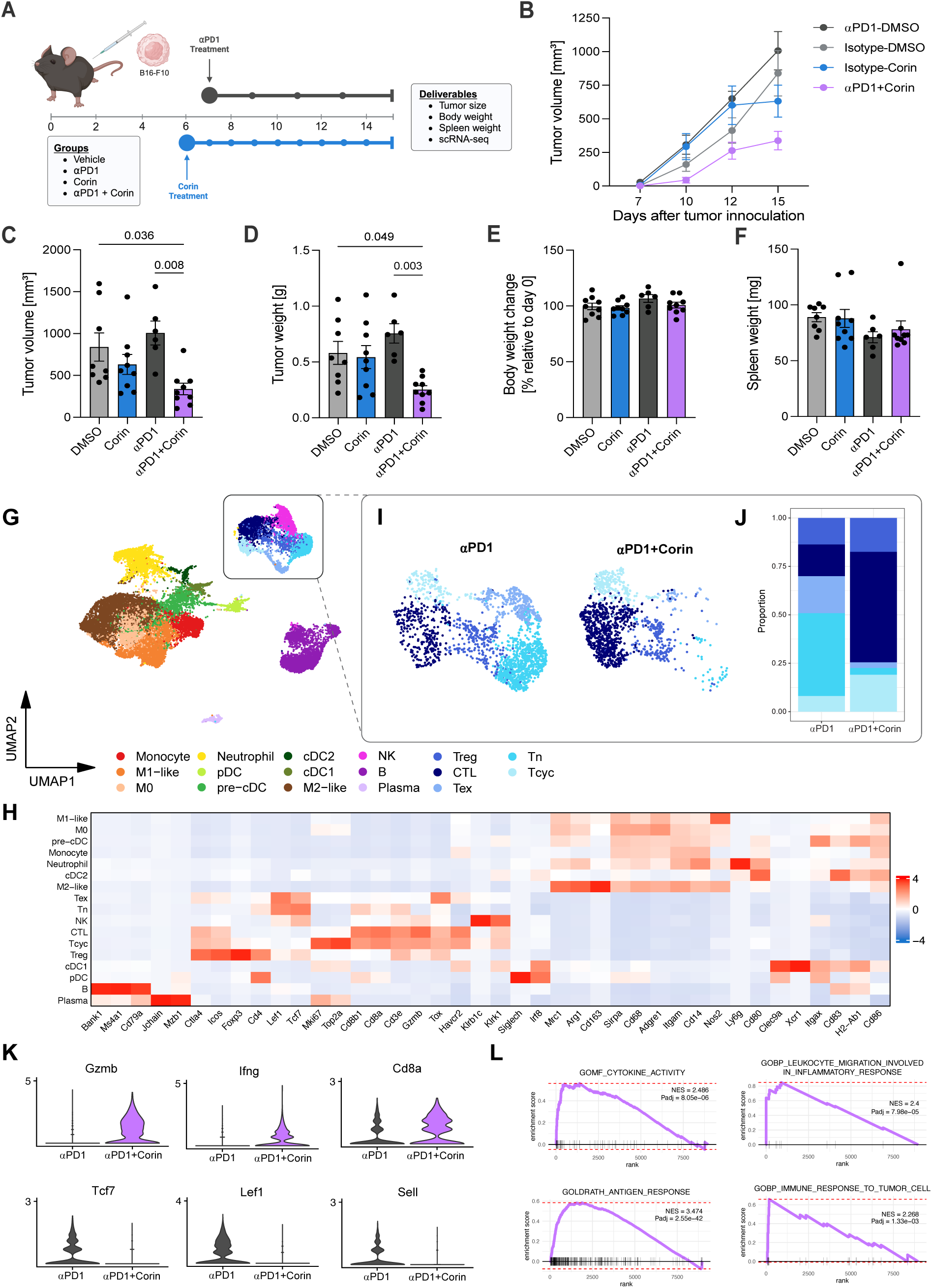
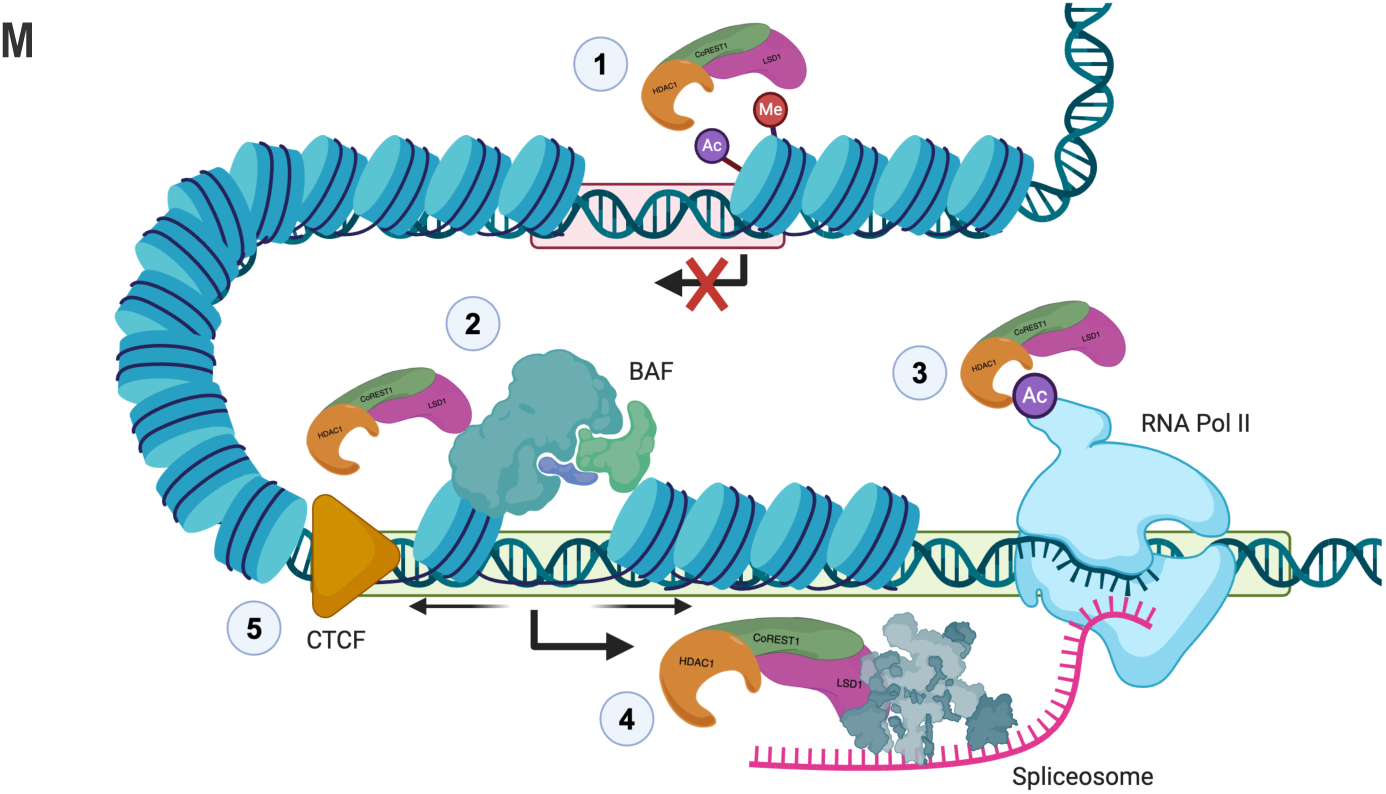
Corin sensitizes immune cold tumors to immunotherapy and promotes expansion of tumor infiltrating cytotoxic T cells. **(A)** Schematic for corin + immunotherapy combination treatment in a melanoma xenograft mouse model. 6-10 week-old female C57BL/6 mice were inoculated with 2.5x10^5^ B16-F10 cells. Mice were treated with 200 µg/mouse of corin or 200 µl vehicle control (5% DMSO/PBS) by daily intraperitoneal injection starting from day 6 after tumor initiation. For anti-PD1 treatment, mice were treated with 150 µg/mice anti-PD1 or isotype control antibody 3 times/week starting from day 7 post-tumor grafting. 10 mice were included in each treatment group. Tumors were measured 3 times/week and tumor volume, tumor weight, body weight change, spleen weight were measured. **(B)** Line plot and **(C)** quantification of tumor volumes from day 7 to day 15 comparing DMSO, ⍺-PD1, corin, and ⍺-PD1 + corin treatment. **(D)** Histogram of tumor volumes depicted in (b). **(E)** Histogram of body weight change relative to day 0 in animals treated with DMSO, ⍺-PD1, corin, and ⍺-PD1 + corin. **(F)** Histogram of spleen weights in animals treated with DMSO, ⍺-PD1, corin, and ⍺-PD1 + corin. Statistical analyses for panels (c-f) were performed using an ordinary one-way ANOVA with Holm-Sidak’s correction for multiple comparisons. Error bars represent the standard deviation (SD). **(G)** scRNA-seq UMAP of the immune population (CD45+) isolated from B16-F10 melanomas. **(H)** Heatmap of the marker genes used to define immune subpopulations in (G). **(I)** Subset UMAP of the T cell compartment comparing ⍺-PD1 treatment to the combination of ⍺-PD1 + corin. **(J)** Stacked barplot of the T cell compartments in (I) **(K)** Violin plots of significant DEGs (Log2FC > |1|, p.adj < 0.05) in T cell populations isolated from ⍺-PD1 versus ⍺-PD1 + corin-treated B16-F10 melanomas. **(L)** GSEA plots for T cell populations isolated from ⍺-PD1 versus ⍺-PD1 + corin-treated B16-F10 melanomas showing enrichment for cytokine activity, leukocyte migration in inflammation, antigen response and immune response in the ⍺-PD1 + corin-treated tumors. **(M)** Schematic illustration of CoREST complex functions and layers of epigenetic crosstalk which ensure transcript fidelity at transcriptionally repressed (red box) and transcriptionally active (green box) regions. (1) Canonical function of the CoREST complex in epigenetic repression through removal of active marks from histone tails. (2) CoREST complex interactions with the nucleosome-remodeling complex, BAF, promote open chromatin structure at transcriptionally active sites. (3) CoREST complex deacetylation of Pol II CTD impacts transcriptional kinetics (4) CoREST complex interactions with the pre-mRNA splicing machinery impacts RNA splicing, alternative transcript expresson and the development of neoantigens. (5) CoREST interactions with CTCF and regulation of higher-order chromatin structure.

## DISCUSSION

Regulation of RNA splicing is critical across all cell types to maintain normal biological functions including cell growth and differentiation^63,64^, while perturbations in splicing are widely seen in cancers^65^. Here we establish a novel role for the CoREST complex in mediating tumorigenesis via direct interactions with the RNA splicing machinery as demonstrated by proteomic analysis, biochemical studies, and cryo-EM structural data. Further, we define a role for corin in inhibiting these interactions and promoting alternative splicing in tumor cells. Remarkably, we find that the CoREST complex modulates pre-mRNA splicing via a variety of mechanisms including direct binding to the 3’ splice site recognition factor, U2AF2, and enhanced splicing factor transcription, both of which are inhibited by corin. Importantly, we define the region of binding of LSD1/RCOR1 to U2AF2 as being within the RRM domain of U2AF2^66,67^, which interfaces with the DNA binding domain of LSD1^47^ suggesting potential competing affinities for LSD1 at this site for transcriptional versus splicing-associated regulatory functions. While we did not include RNA in our cryo-EM sample, the position of the RRM2 domain of U2AF2 could also accommodate bound RNA if it binds to RRM2 in a similar manner as depicted in the AlphaFold prediction. Additionally, we find both LSD1 and RCOR1 as well as splicing factors U2AF2 and SRSF1 to be located at critical splice junctions of tumor-associated splice variants and that these interactions are all inhibited in the setting of corin, further supporting functional interactions of splicing factors and the CoREST complex in cancers.

Our evaluation of CoREST complex effects on pre-mRNA splicing activity allowed us to identify significant corin-induced alternative splicing events across melanomas regardless of tumor subtype, with the greatest influence on skipped exon and retained intron activities. Although such effects of chromatin remodeling enzymes, and HDACs in particular, have been identified in the past and largely been attributable to kinetic coupling of transcription to splicing changes^30^, our PRO-seq analysis of corin-treated cells failed to demonstrate significant changes in Pol II kinetics. This suggests that corin-induced alternative splicing events are the result of direct effects on splicing factor transcription and interactions of the CoREST complex with the splicing machinery. Remarkably, we find that corin-induced splicing changes are not unique to melanoma, as common significant pathways are also seen in ATRT and breast cancers suggesting that corin-induced splicing changes may be more generally applicable to a large number of cancers. Additionally, we demonstrate that splicing events induced by corin lead to the expression of neopeptides that elicit an immune response *in vitro*, while corin treatment of melanoma in an immune competent mouse model leads to significant inhibition of tumor growth in conjunction with immune checkpoint blockade (ICB), rendering an immune-cold melanoma sensitive to checkpoint inhibition. Collectively, these data suggest the potential significant and broad application of corin to enhance immune-mediated responses to cancers.

While previous investigations have demonstrated the utility of pharmacologic induction of RNA splicing to promote neoantigen expression and anti-tumor immunity in preclinical studies^15,16^, and several drugs targeting the splicing machinery have been evaluated in clinical trials^5,68^, the vast majority of these reagents have failed in early-stage studies due to significant toxicities^5,69,70^. Our identification of a novel role for the CoREST complex in binding to splicing factors and the use of the small molecule CoREST inhibitor, corin, to promote alternative splicing events in cancer leading to neoantigen expression and T cell-mediated immunity represents a potential approach to promote immunoreactive neoantigen expression in immune-cold tumors. Indeed, studies suggest a link between tumor mutational burden (TMB), the presence of neoantigens on MHC molecules and responses to ICB therapy in cancers^71–74^ we were therefore interested to see that corin induced thousands of alternative splicing events in both a high-TMB tumor (melanoma) as well as a low-TMB tumor (ATRT) suggesting that corin treatment of ATRT may also result in significant neoantigen expression and induced tumor immunity. We also note that corin induced alternative splicing events at critical splice junctions to a greater extent than either HDACi (MS-275) or LSDi (GSKLSD1) alone or in combination suggesting improved efficacy of corin versus either agent not only on tumor growth^36^, but also within the context of induced splicing changes. Indeed, a previous study found that MS-275 promotes induction of tumor immune editing of tumor neoantigens and tumor immune responses in combination with ICI with ⍺PD-1 therapy in bladder cancer^75^; however, the doses of MS-275 required to elicit these responses were significantly greater than those used for corin in this study. Importantly, we find that CoREST-regulated splicing factors are overexpressed across many tumor types compared to their respective normal tissues, suggesting that cancer cells are more vulnerable to corin-induced splicing changes due to tumor-associated functions of splicing factors. Given the specific targeting of the CoREST complex by corin and the lack of appreciable in vivo toxicity of corin seen in this study and others^36, 37^, we suggest that this work provides a strong preclinical rationale for the use of CoREST inhibitor therapies in combination with ICIs for melanoma and other cancers.

Our finding of direct and functionally significant interactions between CoREST and the splicing machinery represents one of the first reports of direct interactions between a histone modifying complex and the splicing machinery; furthermore, our cryo-EM data represent, to our knowledge, the first reported structure of a histone modifying complex bound to a splicing factor. Given our findings of CoREST complex interactions with the RNA splicing machinery and its specific regulation of transcription and splicing of pre-mRNA splicing factors, along with previous findings of CoREST complex-mediated post-translational modification of RNA-polymerase II^35^, known interactions with transcription factors including SNAG-domain proteins^76^ and REST, as well as the BAF complex^42^ and CTCF, we propose the following model for CoREST complex regulation of transcription that allows for numerous interacting mechanisms to ensure transcript fidelity, which are disrupted in cancer (**Fig. 7M**). We suggest that the CoREST complex plays a critical role in ensuring coordinated regulation of chromatin accessibility, transcription factor binding to relevant DNA sites, transcription kinetics and RNA splicing, and that the CoREST complex provides a means of crosstalk between these systems to ensure proper transcript expression and timing. While we demonstrate the significance of these interactions here in the context of tumorigenesis, we expect these higher order regulatory functions of CoREST, and likely other epigenetic complexes, will prove significant within broader biological processes and provide a critical framework for the tightly coordinated epigenetic control of gene expression.

## RESOURCE AVAILABILITY

### Lead Contact

Further information and requests for resources and reagents should be directed to and will be fulfilled by the lead contact, Rhoda M. Alani.

### Materials Availability

This study did not generate new unique reagents.

### Data and Code Availability

Raw data, processed data, and metadata from the bulk RNA-seq (GSE280449), scRNA-seq (GSE280450), and PRO-seq (GSE280448) experiments have been deposited in the NCBI’s Gene Expression Omnibus (GEO) database. Raw MS data for the CoREST IP-MS are available on PRIDE (project accession: PXD056700, Token: MxX17vDulsT2) and neopeptide IP-MS are available on MassIVE (MHC peptides; MSV000096071). All data supporting the graphs and tables are available upon request. All code used for analysis of sequencing were deposited on GitHub (https://github.com/robertfisher002/CoREST_Splicing; branch main; commit ID 92ac17b). Cryo-EM map and model were deposited in the Protein Data Bank (PDB) and Electron Microscopy Data Bank (EMDB) under the following accession codes: (PDB: 9DWU, EMDB: 47265). Raw cryo-EM movies will be uploaded to the EMPIAR database.

## MATERIALS AND METHODS

### Animal Studies

6-10 week-old female C57BL/6 mice (Jackson Lab) were inoculated with 2.5x10^5^ B16F10 cells. Mice were randomly assigned treatment groups (vehicle control, anti-PD1, corin, and anti-PD1) and treated with 200 µg/mice of corin (HY-111048, MedChemExpress) or 200 µl vehicle control (5% DMSO/PBS) by daily intraperitoneal (i.p.) injections starting from day 6 after the tumor injection. For anti-PD1 treatment, mice were treated with 150 µg/mice anti-PD1 (clone 29F.1A12, BioLegend, 135248) or isotype control (clone RTK2758, BioLegend, 400566) 3 times/week starting from day 7 post-tumor grafts. Treatments occurred blinded and 10 mice per treatment group were used. Tumors were measured 3 times/week and volumes were calculated using the formula V = [1.58π x (length x width)3/2]/6^77^.

### Tissue Processing

Minced tumor biopsies were incubated with 200µL 20mg/mL collagenase, 200µL 20mg/mL hyaluronidase, and 5µL DNase for 20 minutes to 1 hour with agitation until the tumor fully dissociated. Single-cell suspension was passed through a 100µm strainer and washed with 1X PBS by centrifugation. Cells were resuspended in 1ml of freezing media frozen down.

### Cell lines

Melanoma cell lines 451Lu, 1205Lu, SKMEL24, WM793, and SKMEL28 were obtained from Meenard Heryln (The Wistar Institute, Philadelphia, Pennsylvania, USA). SKMEL5 cells were obtained from Deborah Lang (Boston University, Boston, Massachusetts, USA). Cells were maintained at 37C at 5% CO2 and cultured in DMEM (Invitrogen, ThermoFisher Scientific) supplemented with 1% penicillin/streptomycin, L-glutamine (2mM), and 10% FBS. B16F10 cells were cultured in DMEM media (Gibco, 10566024) supplemented with 10% FBS and 100 U/ml of Pen/Strep (Gibco, 15070063). Cells were regularly tested negative for mycoplasma using MycoStrip® -Mycoplasma Detection Kit (InvivoGen, rep-mysnc-50).

### Compounds

Corin (no. HY-111048), MS275 (etinostat, no. HY-12163), and GSK-LSD1 (HY-100546A) were purchased from MedChemExpress. Stock solutions were made in DMSO and an equivalent volume of DMSO was used as a vehicle control.

### Co-Immunoprecipitation

Cells were treated with DMSO or Corin to a final concentration of 2.5µM and incubated for 24 hours. After harvesting in PBS, cell extracts were prepared by resuspending the cell pellet in IP buffer (20 mM Tris [pH 7.5], 137 mM NaCl, 1 mM CaCl₂, 1% NP-40, 10% glycerol, 1 mM MgCl₂, 1X protease and phosphatase inhibitor cocktail; ThermoFisher Scientific, and 0.0125 U/ml Benzonase; Sigma-Aldrich) and rotating at 4°C for 1 hour. Protein concentration was measured using a BCA assay. ProteinA-Dynabeads (Invitrogen, 30 µl per sample) were incubated with antibodies (**Supplementary Table 7**) at 4°C for 1 hour and washed with blocking buffer (0.5% BSA in PBS). 1mg of cell extract were added to antibody-bound beads and incubated overnight at 4°C. After extensive washing with IP buffer lacking MgCl2, protease and phosphatase inhibitor cocktail, and Benzonase, immunoprecipitated proteins were eluted, resolved by SDS-PAGE and subjected to immunoblot analyses.

### S-trap Protein Digestion

Immunoprecipitates from two biological replicates were eluted in a buffer containing 5% SDS, 5 mM DTT and 50 mM ammonium bicarbonate (pH = 8), and left on the bench for about 1 hour for disulfide bond reduction. Samples were then alkylated with 20 mM iodoacetamide in the dark for 30 minutes. Afterward, phosphoric acid was added to the sample at a final concentration of 1.2%. Samples were diluted in six volumes of binding buffer (90% methanol and 10 mM ammonium bicarbonate, pH 8.0). After gentle mixing, the protein solution was loaded to an S-trap filter (Protifi) and spun at 500 g for 30 sec. The sample was washed twice with binding buffer. Finally, 1 µg of sequencing grade trypsin (Promega), diluted in 50 mM ammonium bicarbonate, was added into the S-trap filter and samples were digested at 37°C for 18 h. Peptides were eluted in three steps: (i) 40 µl of 50 mM ammonium bicarbonate, (ii) 40 µl of 0.1% TFA and (iii) 40 µl of 60% acetonitrile and 0.1% TFA. The peptide solution was pooled, spun at 1,000 g for 30 sec and dried in a vacuum centrifuge. Prior to mass spectrometry analysis, samples were desalted using a 96-well plate filter (Orochem) packed with 1 mg of Oasis HLB C-18 resin (Waters). Briefly, the samples were resuspended in 100 µl of 0.1% TFA and loaded onto the HLB resin, which was previously equilibrated using 100 µl of the same buffer. After washing with 100 µl of 0.1% TFA, the samples were eluted with a buffer containing 70 µl of 60% acetonitrile and 0.1% TFA and then dried in a vacuum centrifuge.

### LC-MS/MS Acquisition and Analysis

Samples were resuspended in 10 µl of 0.1% TFA and loaded onto a Dionex RSLC Ultimate 300 (Thermo Scientific), coupled online with an Orbitrap Fusion Lumos (Thermo Scientific). Chromatographic separation was performed with a two-column system, consisting of a C-18 trap cartridge (300 µm ID, 5 mm length) and a picofrit analytical column (75 µm ID, 25 cm length) packed in-house with reversed-phase Repro-Sil Pur C18-AQ 3 µm resin. Peptides were separated using a 90 min gradient from 4-30% buffer B (buffer A: 0.1% formic acid, buffer B: 80% acetonitrile + 0.1% formic acid) at a flow rate of 300 nl/min. The mass spectrometer was set to acquire spectra in a data-dependent acquisition (DDA) mode. Briefly, the full MS scan was set to 300-1200 m/z in the orbitrap with a resolution of 120,000 (at 200 m/z) and an AGC target of 5x10e5. MS/MS was performed in the ion trap using the top speed mode (2 secs), an AGC target of 1x10e4 and an HCD collision energy of 35. Proteome raw files were searched using Proteome Discoverer software (v2.5, Thermo Scientific) using SEQUEST search engine and the SwissProt human database. The search for total proteome included variable modification of N-terminal acetylation, and fixed modification of carbamidomethyl cysteine. Trypsin was specified as the digestive enzyme with up to 2 missed cleavages allowed. Mass tolerance was set to 10 pm for precursor ions and 0.2 Da for product ions. Peptide and protein false discovery rate was set to 1%. Following the search, data was processed as described^78^. Briefly, proteins were log2 transformed, normalized by the average value of each sample and missing values were imputed using a normal distribution 2 standard deviations lower than the mean. Statistical regulation was assessed using heteroscedastic T-test (if p-value < 0.05). Data distribution was assumed to be normal but this was not formally tested.

### Protein purification

The CoREST complex was purified as described (Jay H. Kalin et al., 2018)^36^. Briefly, the constructs of full-length LSD1, full-length HDAC1 and FLAG-tagged NΔ85-CoREST1 were cloned into pcDNA3.0-based vectors and transiently transfected into HEK293F suspension cells using branched polyethylenimine. After 48 hours of incubation, cells were harvested, lysed in cold lysis buffer (50 mM Tris–HCl [pH 7.5], 50 mM KCl, 5% glycerol, 0.4% Triton X-100, 1X Roche EDTA-free Complete Protease Inhibitor cocktail), and sonicated. Insoluble fractions were removed by centrifugation, and the complex was purified via FLAG affinity chromatography. After washing, the complex was cleaved with TEV protease and further purified by Size Exclusion Chromatography using a Superose 6 10/300 GL column (GE Healthcare). For GST-tagged U2AF2 and SRSF1, constructs were cloned into pGEX6P1 vector, expressed in *Escherichia coli*. Cells were harvested and resuspended in GST binding buffer (20mM Tris [pH8.0], 0.2mM EDTA, 1M NaCl, 10% Glycerol, 0.1% NP-40, 1mM PMSF), and lysed using a french press. The lysate was cleared by centrifugation, and the supernatant was incubated with glutathione-Sepharose 4B resin (Cytiva) for 2 hours at 4°C. The resin was extensively washed with GST binding buffer, and the GST-tagged proteins were eluted with GST binding buffer containing 30 mM glutathione. The eluted proteins were dialyzed against dialysis buffer (20mM Tris [pH8.0], 0.2mM EDTA, 150mM NaCl, 10% Glycerol), then stored at -80°C. For His-tagged fl-LSD1, truncated LSD1/RCOR1 complex^79^ and U2AF2 2 a.a. 41-471, constructs were cloned into a pET15b vector (His-tagged fl-LSD1, truncated His-tagged LSD1, and U2AF2) and pET-28b (His-tagged truncated RCOR1). The plasmid encoding His-tagged fl-LSD1 was a kind gift from Dr. Yang Shi (University of Oxford). For the purification of fl-LSD1, truncated LSD1/RCOR1 complex and U2AF2 (aa241-471), His binding buffer (20mM Tris [pH 8.0], 300mM NaCl, 1mM TCEP) was used for cell resuspension, while the resin was washed with His binding buffer containing 40 mM imidazole, and elution was performed using His binding buffer containing 200 mM imidazole. The eluants were dialyzed against the His binding buffer containing 1 mM EDTA. For His-LSD1, additional purification was performed using MonoQ (Cytiva) and Superdex 200 increase 10/300 GL columns (Cytiva).

### *In vitro* pull-down assay

GST-tagged splicing factors (final concentration 300 nM) were mixed with the CoREST complex or LSD1 (final concentration 15 nM), 15 µl of glutathione-Sepharose 4B resin (Cytiva), and BSA (final concentration 0.3 mg/ml) in binding buffer (20 mM Tris [pH 8.0], 0.2 mM EDTA, 150 mM NaCl, 10% glycerol, 0.1% NP-40, 1 mM PMSF) to a total volume of 300 µl. The mixtures were incubated with rotation at 4°C for 2 hours, after which the resin was washed four times with binding buffer. The resin-bound proteins were eluted, resolved by SDS-PAGE and subjected to immunoblot analyses.

### Western blot

Whole-cell protein lysates were extracted in MPER buffer supplemented with Halt Protease Inhibitor. Samples were run on 10% SDS-PAGE gels and transferred to polyvinylidene difluoride membranes. Membranes were blocked with 5% nonfat dry milk diluted in a 0.05% Tween 20 PBS solution and incubated overnight at 4C with primary antibody (**Supplementary Table 7**). Membranes were washed 3x with Tris Buffered Saline with Tween 20 (TBST) then incubated with HRP-conjugated secondary antibodies (**Supplementary Table 7**) for 1 hour, then washed again 3x with TBST before visualized with ECL Western Blot Substrate (ThermoFisher Scientific). Chemiluminescent blots were imaged with the Chemi-doc XP Imager (Bio-Rad) and analyzed using ImageJ densitometry quantification. All bands were normalized against GAPDH for quantification.

### Cryo-EM sample preparation

Frozen samples of truncated LSD1/RCOR1 complex (6xHis-LSD1 (aa171-852) + 6xHis-RCOR1 (aa289-485)) and 6xHis-U2AF2 (aa241-475) were thawed on ice, mixed at a molar ratio of 1:5. The protein mixture was dialyzed overnight at 4°C against a dialysis buffer (20 mM Tris [pH 8.0], 150 mM NaCl, and 1 mM TCEP). Following dialysis, the sample was separated using a Superdex 200 Increase 10/300 GL column (GE Healthcare). The peaks were analyzed by SDS-PAGE, and fractions containing LSD1, RCOR1, and U2AF2 were pooled and concentrated for grid freezing. Ultrafoil R1.2/1.3 Au 300 mesh grids were glow-discharged for 45 seconds at 15mA using a PELCO easiGLOW Glow Discharge System to apply a negative charge to their surface. Then, 3 μL of sample mixture at a concentration of 0.75 mg/mL was applied to the grid, immediately blotted for 3 seconds with a blot force of 3, and plunge-frozen in liquid ethane using a Vitrobot Mark IV apparatus (Thermo Fisher) set to 100% humidity and 4°C.

### Cryo-EM data collection

A cryo-EM dataset of truncated LSD1/RCOR1 complex bound to U2AF2 was collected at the Boston University Cryo-EM Core Facility using a 200 kV ThermoFisher Glacios 2 cryoEM microscope equipped with a Falcon 4i direct electron detector and Selectris energy filter. A dataset of 12,845 exposures was collected in counting mode and recorded in Electron Event Representation (EER) format using a magnification of 130kx, pixel size of 0.90 Å, nominal dose of 50 e-/ Å^2^, dose rate of 10.84 e-/px/s, a defocus range of -1.0 to -2.5 μm, and an energy filter slit width of 10 eV. A multi-shot imaging strategy was used to collect 3 shots per hole, utilizing beam image shift to move between each target.

### Cryo-EM data processing

The cryoEM dataset was processed in cryoSPARC v4.5.1^80^. Exposures were imported with an EER upsampling factor of 2 and cropped to one-half their original resolution using Patch Motion Correction. The CTF was corrected using Patch CTF Estimation, and poor-quality micrographs were removed yielding 6,118 high-quality micrographs. An initial set of blob picks from a small subset of micrographs was classified using 2D Classification to create a set of high quality 2D templates. The 2D templates were used to perform particle picking using Template Picker and Inspect Picks, then extracted at a box size of 360 pixels using Extract from Micrographs to yield an uncleaned particle stack (3,973,676 particles). Two rounds of 2D Classification with a circular mask of 150 Å, initial classification uncertainty factor of 4, number of online-EM iterations of 30, and Batchsize per class of 300, were performed to remove junk particles by discarding poor-quality 2D classes to yield a partially cleaned particle stack (1,183,288 particles). Two rounds of parallel multi-structure Ab-initio reconstruction jobs were performed where particles from distinctly good volumes were selected yielding a cleaned particle stack (961,750 particles). Homogenous Refinement with Adaptive Marginalization, Non-Uniform Refinement^81^, and Local Refinement with pose/shift gaussian prior were performed consecutively to align the cleaned particle stack. Rebalance Orientations was performed on the aligned cleaned particle stack to remove particles from oversampled preferred orientations, yielding a balanced aligned cleaned particle stack (695,785 particles). Focused 3D classification at a target resolution of 15 Å and 4 classes was performed using a spherical focus mask centered on the density corresponding to U2AF2. Homogenous Reconstruction Only was performed on all four particle stack corresponding to the 3D volume outputs. The reconstruction with the strongest U2AF2 density was selected as the final reconstruction of truncated LSD1/RCOR1 complex bound to U2AF2 (172,429 particles, 5.14 Å).

### Cryo-EM model building

Initial model of truncated LSD1/RCOR1 complex bound to U2AF2 was built by rigid body fitting models of LSD1+CoREST (PDB: 2IW5)^46^ and U2AF2 (PDB: 5W0H)^66^ into the EM density map using ChimeraX^82^. Hydrogen atoms and alternate side-chain rotamers were removed from the initial model using PyMOL. To account for missing EM density in the region of the RCOR1 protein, residues 377-440 were removed from RCOR1. Individual Ramachandran and rotamer outliers were corrected where applicable using Coot 0.9.6^83^, and the final model was validated using cryo-EM Comprehensive Validation module in PHENIX^84^ running MolProbity^85^. Figures were generated using ChimeraX^82^.

### RNA-sequencing

Total RNA was extracted from melanoma cell lines treated with corin or vehicle control (DMSO) using the Qiagen RNeasy Mini Kit. Samples were quantified using nanodrop and sequenced by Azenta Inc.

### RNA-seq data analysis

Paired-end RNA-seq reads (2x150bp) were quality and adaptor-trimmed using Trimmomatic (v.0.36)^86^. Trimmed reads were mapped to the ENSEMBL reference human genome (GRCh38) using STAR (v.2.5.2b)^87^. BAM files were generated for downstream splicing and differential gene expression analysis. Unique gene counts were calculated using featureCounts from Subread (v.1.5.2)^88^ and differential expression analysis was performed using DESeq2 (v.1.44.0)^89^ based on hit counts. The Wald test was used to produce log2 fold change and p-values between comparisons. A threshold of p < 0.01 and LF > |0.5| was used to call significant changes in gene expression. Gene set enrichment analysis was performed using GSEA (v.4.3.2)^90^ with KEGG^91^ and Hallmark^92^ curated gene sets.

### Melanoma splicing analysis

Two tools were used to identify RNA splicing changes from RNA-seq read data. rMATs-turbo (v.4.2.0)^52^ was used to call skipped exon, alternative 3’SS, alternative 5’SS, and mutually exclusive events, while LeafCutter (v.0.2.9)^53^ was used to call intron retention events. rMATS was run using default parameters for BAM file inputs and the --allow-clipping parameter to prevent soft-clip skipping. LeafCutter was run using default parameters for BAM file inputs. A threshold of q < 0.05 and deltaPSI ≥ |0.1| was used to call significant differential events. Gene ontology analysis and visualization was performed on significant SE events using the R package enrichplot (v.1.22.0). Sashimi plots were generated using rmats2sashimiplot (v.2.0.4).

### ATRT and breast cancer splicing analysis

Corin-treated breast cancer FASTQ files were downloaded with SRAtoolkit from GEO (Series GSE168644). BAM files were generated using STAR (2.5.2b)^87^ against the hg38 reference genome. ATRT FASTQ samples were acquired from collaborators at Johns Hopkins University and aligned to the hg38 reference genome using STAR (2.5.2b)^87^. Differential splicing analysis was conducted using rMATS-turbo (v.4.2.0)^52^ and LeafCutter (v.0.2.9)^53^. Significant events were called using q < 0.05 and |deltaPSI| ≥ 0.1.

### TCGA splicing data analysis

OncoSplicing^57^ was used to identify survival-associated splicing events (p <0.05) in TCGA-SKCM. Hazard ratios and Kaplan-Meier plots were stored if an event appeared in the list of significant corin-induced splicing events and in the OncoSplicing database. cBioPortal^93^ was used to identify the alteration frequency of corin-affected RBP expression across cancers with matched normal tissue data (TCGA pancancer dataset).

### RT-PCR splice gels

Total RNA was extracted using the Qiagen RNeasy Mini Kit. cDNA was synthesized with the SuperScript cDNA Synthesis Kit (ThermoFisher Scientific) and splice products were amplified with splice site-specific primers (**Supplementary Table 8**) and Taq polymerase for 35 cycles at Tm = 52.5C. DNA was run on 1.5% agarose gels and quantified using inclusion:exclusion product ratios.

### RNA immunoprecipitation coupled with quantitative PCR (RIP qPCR)

SKMEL5 cells were treated with DMSO or Corin to a final concentration of 2.5µM and incubated for 24 hours. Cells were harvested PBS, lysed in RNA lysis buffer (25 mM Tris [pH 7.5], 150 mM NaCl, 5 mM EDTA, 5 mM MgCl2, 1% NP-40, 0.5 mM DTT, 1X protease and phosphatase inhibitor cocktail; ThermoFisher Scientific, 40 U/ml RNase inhibitor; Applied Biosystems), and incubated for 30 minutes at 4°C with rotation. Chromatin was sheared by passing through a 22-gauge needle, and lysates were centrifuged. The supernatant was mixed with 1.5 mg of cell extract and specific antibodies (**Supplementary Table 7**) and incubated for 2 hours at 4°C. ProteinA Dynabeads (Invitrogen, 30 µl per sample) were washed and added to each sample for 2.5 hours, followed by three washes with RNA lysis buffer. The beads were then treated with TRIzol for RNA extraction. RNA was purified by chloroform extraction and isopropanol precipitation, followed by DNase treatment and cDNA synthesis using the random hexamer and SuperScript cDNA Synthesis Kit (ThermoFisher Scientific). Quantitative PCR (qPCR) was performed using SYBR^®^ Green Quantitative RT-qPCR kit (Sigma Aldrich) and splice site-specific primers (**Supplementary Table 8**). The data analysis was performed by calculating ΔCq normalized to β-actin expression by the StepOnePlus.

### PRO-seq library construction

SKMEL5 cells were treated for 24h with 2.5µM corin or DMSO and permeabilized as described. All sample preparation was conducted on ice (4°C). Cells were washed in ice cold 1x PBS and resuspended in wash buffer (10 mM Tris-HCl pH 8.0, 10% glycerol, 250 mM sucrose, 10 mM KCl, 5 mM MgCl2, 0.5 mM DTT, 1mM EGTA, Halt protease inhibitor cocktail (Thermo Scientific), and 4 u/mL RNase inhibitor [SUPERaseIN, Invitrogen]). Then cells (2x10^7) were gently permeabilized in permeabilization buffer (10 mM Tris-HCl pH 8.0, 10% glycerol, 250 mM sucrose, 10 mM KCl, 5 mM MgCl2, 0.5 mM DTT, 0.1% Igepal, protease inhibitors cocktail (Roche), 4µ/mL RNase inhibitor [SUPERaseIN, Invitrogen]) for 5 minutes. Cells were recovered by centrifugation (400 x g for 8 minutes) and the supernatant was carefully removed. Cells were washed with 10 mL of wash buffer and then centrifuged again under the same conditions. Finally, cells were resuspended in 400µL of freeze buffer (50 mM Tris-HCl pH 8.0, 40% glycerol, 5 mM MgCl2, 0.5 mM DTT, 4 u/mL RNase inhibitor [SUPERaseIN, Invitrogen]) and stored at -80°C.

Aliquots of frozen (-80°C) permeabilized cells were thawed on ice and pipetted gently to fully resuspend. Aliquots were removed and permeabilized cells were counted using a Luna II, Logos Biosystems instrument. For each sample, 1 million permeabilized cells were used for nuclear run-on, with 50,000 permeabilized *Drosophila* S2 cells added to each sample for normalization. Nuclear run on assays and library preparation were performed essentially as described in Reimer et al. [K. A. Reimer, C. A. Mimoso, K. Adelman, K. M. Neugebauer, Molecular Cell (2021)] with modifications noted: 2X nuclear run-on buffer consisted of (10 mM Tris (pH 8), 10 mM MgCl2, 1 mM DTT, 300mM KCl, 20uM/ea biotin-11-NTPs (Perkin Elmer), 0.8U/µl SuperaseIN (Thermo), 1% sarkosyl). Run-on reactions were performed at 37°C. Random hexamer extensions (UMIs) were added to the 3’ end of the 5’ adapter and 5’ end of the 3’ adapter. Adenylated 3’ adapter was prepared using the 5’ DNA adenylation kit (NEB) and ligated using T4 RNA ligase 2, truncated KQ (NEB, per manufacturer’s instructions with 15% PEG-8000 final) and incubated at 16°C overnight. 180µl of betaine buffer (1.42g of betaine brought to 10mL) was mixed with ligations and incubated 5 min at 65°C and 2 min on ice prior to addition of streptavidin beads. After T4 polynucleotide kinase (NEB) treatment, beads were washed once each with high salt, low salt, and 0.25X T4 RNA ligase buffer (NEB) and resuspended in 5’ adapter mix (10 pmol 5’ adapter, 30 pmol blocking oligo, water). 5’ adapter ligation was per Reimer but with 15% PEG-8000 final. Eluted cDNA was amplified 5-cycles (NEBNext Ultra II Q5 master mix (NEB) with Illumina TruSeq PCR primers RP-1 and RPI-X) following the manufacturer’s suggested cycling protocol for library construction. A portion of preCR was serially diluted and for test amplification to determine optimal amplification of final libraries. Pooled libraries were sequenced using the Illumina NovaSeq platform

### PRO-sequencing data analysis

All custom scripts described herein are available on the AdelmanLab GitHub (https://github.com/AdelmanLab/NIH_scripts). Dual, 6nt Unique Molecular Identifiers (UMIs) were extracted from read pairs using UMI-tools [10.1101/gr.209601.116]. Read pairs were trimmed using cutadapt 1.14 to remove adapter sequences (-O 1 --match-read-wildcards -m {20,26}). The UMI length was trimmed off the end of both reads to prevent read-through into the mate’s UMI, which will happen for shorter fragments. An additional nucleotide was removed from the end of read 1 (R1), using seqtk trimfq (https://github.com/lh3/seqtk), to preserve a single mate orientation during alignment. The paired end reads were then mapped to a combined genome index, including both the spike (dm6) and primary (hg38) genomes, using bowtie2 [10.1038/nmeth.1923]. Properly paired reads were retained. These read pairs were then separated based on the genome (i.e. spike-in vs primary) to which they mapped, and both these spike and primary reads were independently deduplicated, again using UMI-tools. Paired-end RNA-seq reads were mapped to the hg38 reference genome via HISAT2 v2.2.1 (--known-splicesite-infile). To select gene-level features for differential expression analysis, and for pairing with PRO-seq data, we assigned a single, dominant TSS and transcription end site (TES) to each active gene. This was accomplished using a custom script, get_gene_annotations.sh (available at https://github.com/AdelmanLab/GetGeneAnnotation_GGA), which uses RNA-seq read abundance and PRO-seq R2 reads (RNA 5’ ends) to identify dominant TSSs, and RNA-seq profiles to define most commonly used TESs. RNA-seq and PRO-seq data from all conditions were used for this analysis, to comprehensively capture gene activity in these samples.Reads were summed within the TSS to TES window for each active gene using the make_heatmap script (https://github.com/AdelmanLab/NIH_scripts), which counts each read one time, at the exact 3’ end location of the nascent RNA. DEseq2, using the Wald test, was used to determine statistically significant differentially expressed genes. Unless otherwise noted, the default size factors determined by DEseq2 were used.

### Neopeptide prediction analysis

Two pipelines were used to predict splice-induced neopeptide candidates. First, JCEC files containing significant rMATS SE events (q < 0.05, deltaPSI ≥ |0.1|, *log*2(TPM) ≥ 3, IJC + SJC ≥ 20) were input into SpliceTools^58^ to translate and identify peptide sequences produced by frameshifted events. Then, sequences were digested into unique 8-11mer sequences and filtered against the human proteome (UCSC) to isolate neopeptide products. Second, the SNAF^54^ T-cell pipeline with default parameters was used to capture additional neopeptides produced from RI, A5’SS, A3’SS, MXE, and trans-splicing events. Predicted neopeptides were validated against the human proteome and appended to the SpliceTools-identified events. Then, MHC binding predictions were performed using both HLAthena^60^ and NetMHCpan (v.4.1)^59^ with the following parameters “-l 8, 9, 10, 11” and %Rank ≤ 2. Neopeptides identified by both prediction tools were selected and ranked based on a score defined by the product of the -*log*10(%Rank), junction count, and |deltaPSI| value.

### MHC I Immunoprecipitation, peptide purification, and mass spectrometry

Immunoprecipitation with W6/32 MHC Class I antibody (Santa Cruz Biotechnology #SC-32235) and LC-MS/MS analysis were performed following previously established protocols (Klaeger et al., 2021; Sarkizova et al., 2020). HLA peptides were extracted by reconstituting washed beads in 3% acetonitrile/5% formic acid and shaking for 3 minutes. Beads and supernatant were transferred to spin columns (Pierce). Original tubes were washed 2x with 0.1% formic acid and washings added to the spin column. Spin column elutions were applied to the well of a previously activated (80% acetonitrile, 0.1% formic acid) and equilibrated (0.1% formic acid) 40 mg tC18 plate and vacuum applied to pull liquid through. Beads (still on the spin column) were further eluted 3x with 10% acetic acid for 3 minutes with shaking. Each time, after diluting to 1% acetic acid, spin column elutions were applied to the same well of the tC18 plate and vacuum applied. The well was then washed 3x with 0.1% formic acid. Desalted peptides were eluted with 15% acetontrile/1% formic acid, then with 50% acetonitrile, 1% formic acid (into the same tube) and dried by vacuum centrifugation. Peptides were reconstituted in 3% acetonitrile, 0.5% formic acid and applied to EVO tips prepared according to the manufacturer’s instructions. EVO tips were eluted twice with 25 µL of 35% acetonitrile/0.1% formic acid, dried by vacuum centrifugation, and stored at -80 °C. For analysis, peptides were reconstituted in 3% acetronitrile/0.5% formic acid, transferred to total recovery vials (Waters, Milford, MA) and analyzed by nanoLC-MS using a NanoElute HPLC system (column=25 cm x 75 µm I.D. packed with 1.9 µm Dr. Maisch C18 with integrated emitted tip^94^ interfaced to a timsTOF SCP mass spectrometer (Bruker, Billerica, MA). Peptides were eluted with an HPLC gradient (0-15% B in 60 minutes, 15-23% B in 90 minutes, 23-35% B in 10 minutes, and 35-80% B in 10 minutes; A=0.1% formic acid, B=0.1% formic acid in acetonitrile) at a flow rate of 200 nL/min. Peptides were analyzed by 10 cycles of DDA-PASEF using a 200 ms ramp from 1/k0 0.6-1.6 and *m/z* 200-1700. Active exclusion was enabled with a release time of 0.4 minutes, threshold was 750 counts, and target was 20000 counts. Peptides derived from SE events and identified by SpliceTools were searched using MSFragger^95^ version 20 installed on Bridges2 (Pittsburgh Supercomputing Facility). Additionally, peptides derived from RI, 3’SS, 5’SS, MXE, or trans-splicing events and identified by the SNAF^54^ pipeline were searched using SNAF’s MaxQuant^96^ wrapper function following default parameters. Peptides with FDR < 0.05 from either analysis were considered. Peptides were classified as neopeptides if the peptide sequence did not appear in the UCSC database containing the human proteome appended with virus and reverse decoy sequences.

### Candidate neoantigen identification

Candidate neoantigens were defined based on two methods: 1) a neopeptide was predicted to bind to human MHC and was identified in at least one corin-treated replicate of mass spectrometry (FDR < 0.05) but not in either DMSO-treated replicate 2) a neopeptide was predicted to bind human MHC, had a high rank score (based on deltaPSI, junction count, and binding %Rank, and a high immunogenicity score from DeepImmuno^61^ but was not recovered by mass spectrometry as previously described^15,16^. Note: C03:04 does not have adequate training data to accurately predict the immunogenicity of bound peptides so only highly ranked A11:01 and B40:01 were tested.

### Peptide synthesis

Candidate neopeptides (10mg) were synthesized by RS Synthesis for ELISpot assays (Louisville, KY, USA) and shown to have > 90% purity by HPLC.

### ELISpot assay

To expand reactive T cells, HLA-matched PBMCs were pre-stimulated with synthesized peptides (10µg/ml) for 14 days in IL2/IL7 media. APCs were isolated from CD4 and CD8 depleted PBMCs, loaded with peptides (10µg/ml), and seeded at a ratio of 3:1 with the pre-stimulated T cells in a 96 well ELISpot plate. IFN-y ELISpot was performed per manufacturer’s instructions and spots were quantified using an Immunospot Analyzer.

### scRNA-seq sample preparation

Cryopreserved single-suspension tumor samples were thawed and dead cells were removed using the Dead Cell Removal Kit (Miltenyi, 130-090-101) following manufacturer instructions. Live cells from the flow-though were used for CD45+ cell isolation using CD45 mouse MicroBeads (Miltenyi, 130-052-301) following the recommendations. CD45+ cells were counted and resuspended in PBS + 0.04% BSA at 1000 cells/µL for scRNA-seq.

### 10x scRNA-seq

About 23,000 cells were loaded onto a 10× Genomics ChromiumTM X instrument (10x Genomics) according to the manufacturer’s recommendations. The scRNAseq libraries were processed using Chromium GEM-X Single Cell 5’ Kit v3 (10x Genomics). Quality controls for amplified cDNA libraries and final sequencing libraries were performed using Bioanalyzer High Sensitivity DNA Kit (Agilent). The sequencing libraries for scRNAseq were normalized to 4nM concentration and pooled. The pooled sequencing libraries were sequenced on the Illumina NovaSeq S4 300 cycle platform. The sequencing parameters were: Read 1 of 28bp, Read 2 of 90bp, Index 1 of 10bp and Index 2 of 10bp.

### Single-cell data processing and cluster annotation

Transcriptomic data were mapped to the Mus musculus reference genome (GRCm39) and assigned to individual cells of origin using Cell Ranger software 8.0 (10X Genomics). Subsequent count matrices were read into Seurat (v5.0.11)^97^ for downstream analysis. Doublets were identified and removed for each sample individually using the DoubletFinder (v2.0.4)^98^ pipeline. The data were further filtered to exclude cells with high mitochondrial gene expression (>10%) and aberrant unique feature counts (<600, >4000) or contaminating non-immune cells. Samples were then merged together for normalization and identification of highly variable features using scTransform (v0.4.13)^99^. Dimensionality reduction was carried out via principal component analysis using 20 dimensions. The optimal number of clusters was identified using clustree (v0.5.1)^100^ and each cluster was annotated based on the top genes determined by FindAllMarkers function in Seurat and SingleR^62^ annotations. The T cell population was further subset based on Cd3e expression to determine effects of corin treatment.

### Single cell differential gene expression and pathway enrichment

Differential expression analysis was conducted via the FindMarkers function in Seurat, using the statistical framework introduced by MAST^101^. To further explore the biological processes and pathways associated with these differentially expressed genes, gene set enrichment analysis was performed using the FGSEA (v1.26.0)^102^ package with 50,000 permutations, drawing on gene sets from the Molecular Signatures Database (MSigDB).

### Single cell differential cell type proportion analysis

A binomial generalized linear model (GLM) framework was employed using the lmerTest (v3.1.3)^103^ package to compare the abundance of different cell types between experimental conditions. Differential cell type proportions were quantified as log odds ratios, indicating enrichment or depletion of cell types across conditions. Estimated marginal means and pairwise contrasts were obtained using the emmeans (v1.10.1) package.

### RBP activity prediction

RBP activity prediction was performed using RNA-SPRINT from the SNAF^54^ pipeline with default parameters. RBP activity scores and matched RBP gene expression (TPM) for each sample were used to calculate Spearman’s statistic using the stats R package (v.3.6.2). U2AF2 motif enrichment coverage plots were generated for significant SKMEL5 SE events using rMAPS2^104^.

### Quantification and statistical analysis

All raw data, source code, and processed data can be found on Github (https://github.com/robertfisher002/CoREST_Splicing/tree/main). Raw sequencing data can be found on NCBI’s GEO.

## Acknowledgments

The authors would like to thank the Nascent Transcriptomics Core at Harvard Medical School, Boston, MA for performing PRO-seq library construction and for assistance with data analysis, Marianne Collard and Ana Fiszbein for helpful discussions and advice, and members of the BU Department of Dermatology for their helpful suggestions and critical review of this work. The authors would like to thank Dr. Yang Shi (University of Oxford) for generously providing the plasmid encoding His-tagged fl-LSD1. This research was supported by the following: Department of Defense Grant W81XWH2110980 (RMA, KP); Melanoma Research Alliance Grant #1045461 (RMA, RJF); American Heart Association (Postdoctoral Fellowship Award 826614 to K.L.); The Hevolution Foundation (AFAR), the Einstein-Mount Sinai Diabetes Center, and the NIH Office of the Director (S10OD030286) (SS); U01 CA243004 and T32 AI007309-33 (CSE); The Swiss National Science Foundation (Grant 225660) and NIH 1R01CA279391-01A1 (KP); National Institutes of Health grant (R01 CA233800) and the Masschusetts Life Science Center (JAM); National Institutes of Health grant (GM149229) (PAC); National Institutes of Health grants (U54-CA272688, RO1-HL157174, RO1-CA285308 & RO1-CA279391) (DBK); This work was supported in part by NIH U24CA224331 (to CJW). CJW is the Lavine Family Chair for Preventative Cancer Therapies. Images were created with BioRender.

## Author contributions

Conceptualization: RMA, RJF

Methodology: RMA, RJF, KP, KL, KP, AV, CE, SF, JM, JC, CWH, DBK, SS, CJW, PAC

Investigation: RJF, KP, KL, KP, AG, ER, CWH, SS

Visualization: RJF, KP, CWH, SS

Funding acquisition: RMA, PAC

Project administration: RMA

Supervision: RMA, PAC, CJW, SS, DBK

Writing – original draft: RMA, RJF

Writing – review & editing: RJF, KP, KL, KP, AV, CE, SF, JM, JC, AG, ER, CWH, DBK, SS, CJW, PAC, RMA

## Competing interests

PAC is a founder of Acylin Therapeutics and has been a consultant for Abbvie, Constellation and Epizyme. He is an inventor of an issued U.S. patent for Corin. JAM is a founder, equity holder, and advisor to Entact Bio, serves on the SAB of 908 Devices, and receives or has received sponsored research funding from Vertex, AstraZeneca, Taiho, Springworks, TUO Therapeutics, and Takeda. DBK is a scientific advisor for Immunitrack, a wholly owned subsidiary of Eli Lilly and Company and Breakbio. DBK owns equity in Affimed N.V., Agenus, Armata Pharmaceuticals, Breakbio, BioMarin Pharmaceutical, Celldex Therapeutics, Editas Medicine, Gilead Sciences, Immunitybio, Lexicon Pharmaceuticals. RMA is a co-founder of Acylin Therapeutics. CJW holds equity at BioNTech, and receives research funding from Pharmacyclics. She is a SAB member of Repertoire, Adventris and Aethon Therapeutics.

---

**Supplementary Fig. 1. Data processing pipeline for cryo-EM analysis and structural superimpositions. (A)** Data processing pipeline to determine the cryo-EM structure of U2AF2 bound to the LSD1+RCOR1 complex. **(B)** Fourier Shell correlation plot showing a global resolution of 5.14 Å. **(C)** Local resolution estimation showing a cryo-EM map resolution range from 5.0 Å to 7.0 Å. **(D)** Superimposition of LSD1+RCOR1 over the EM density showing an excellent fit. **(E)** Superimposition of the RRM2 domain of U2AF2 over the U2AF2 EM density showing a good fit. **(F)** Superimposition of the UHM domain of U2AF2 over the U2AF2 EM density showing a poor fit.

**Supplementary Fig. 2. CoREST inhibition does not impact splicing factor transcript steady state. (A)** Diagram of KEGG Spliceosome pathway with genes downregulated by corin treatment highlighted in pink. **(B)** Western blot biological replicates used in the quantification of Fig. 3E. **(C)** Gene set enrichment analysis for the KEGG Spliceosome gene set using expression data derived from PRO-seq (q < 0.05). **(D)** Heatmap comparing PRO-seq to RNA-seq Log2FC values across the significant downregulated splicing factors.

**Supplementary Fig. 3. CoREST inhibition impacts RNA splicing factor intron retention and selectively skips short exons. (A)** Exon and proximal intron length bias between differentially included and excluded exons. Statistical analysis was performed using ANOVA to compare lengths across treatment groups, grouped by type of splicing event. P-values were adjusted for multiple comparisons using the Holm-Sidak’s correction. **(B)** Exon and proximal GC bias between differentially included and excluded exons. Statistical analysis was performed using ANOVA to compare GC content across treatment groups, grouped by type of splicing event. P-values were adjusted for multiple comparisons using the Holm-Sidak’s correction. **(C)** Percent Spliced In (PSI) distribution across 6 melanoma cells lines for significant Intron Retention (RI) events (deltaPSI ≥ |0.1|, q < 0.05). Statistical comparisons were performed using a two-sample t-test to assess differences in PSI value between treatment groups within each cell line. P-values were adjusted for multiple comparisons using the Bonferroni correction. **(D)** Pathway analysis for significant RI events across the 6 melanoma cell lines (p < 0.05). Enrichment analysis was performed using the hypergeometric test with multiple test correction by the Benjamini-Hochberg method.

**Supplementary Fig. 4. RNA-SPRINT analysis of RBP activity correlated with corin-induced expression changes. (A)** Rank plot of Spearman’s rank correlation between RBP activity and expression with corin treatment. Red points denote RBPs with a correlation value > 0.5. **(B)** RBP activity scores for each candidate RBP determined in (A) across all 6 melanoma cell lines. **(C)** Individual correlation plots for each candidate RBP. **(D)** rMAPS2 coverage plots for U2AF2 binding scores in significantly included (blue) and excluded (red) exons in SKMEL5 cells treated with corin compared to a randomized background list of exons (black). Motif scores are represented by solid lines and p-values are represented by dashed lines. The window is centered at the expons of interest (green box).

**Supplementary Fig. 5. Validation of corin-induced RNA splicing changes. (A)** Sashimi plots of MYO1B, TJP1, and FN1 splicing based on RNA-seq data. **(B)** TJP1 exon 20 RT-PCR gel comparing DMSO, LSD1i, HDACi, LSD1i + HDACi, and corin’s impact on exon inclusion. Statistical analysis of biological replicates (n=3) was performed using an unpaired, two-tailed t-test. Error bars represent the standard deviation (SD). **(C)** RT-PCR gel biological replicates used for quantification in Fig. 4F and S5B.

**Supplementary Fig. 6. Corin promotes promoter pause release without impacting transcriptional kinetics at splice sites**. **(A)** PRO-seq coverage metagene plot centered at TSS’s under DMSO or corin conditions. **(B)** Pausing index calculated between treatments. Statisical analysis performed using a two-sided two-sample t-test. **(C)** PRO-seq coverage metagene plots at 3’SS and 5’SS for significantly included and excluded skipped exon events (deltaPSI ≥ |0.1|, q < 0.05).

**Supplementary Fig. 7. Additional immunological analyses to validate neoantigen immunogenicity and corin’s impact on the tumor microenvironmentS.** ELISpot images and quantification for HLA B40:01 **(A)** and HLA C03:04 **(B)**. Statistical analysis was performed using multiple two-tailed unpaired t-tests. Error bars represent the standard deviation (SD). No peptides reached statistical significance. **(C)** scRNA-seq UMAP comparing the immune populations between anti-PD1 and anti-PD1+corin treatment. **(D)** Proportion analysis comparing changes in the immune populations between treatments. **(E)** Barplots and statistical analysis comparing changes in T cell proportions (*p.adj < 0.05, **p.adj < 0.01, ***p.adj < 0.001). **(F)** Volcano plot of differential gene expression in the T cell subset.

**Supplementary Table 1. Results from the IP-MS using the LSD1 and RCOR1 pulldowns.**

Log2(Fold change) and -Log2(p) values were calculated by comparing signal to IgG control.

**Supplementary Table 2. Results from the IP-MS using the LSD1 and RCOR1 pulldowns.** Log2(Fold change) and -Log2(p) values calculated by comparing DMSO signal to corin signal (24h, 2.5uM).

**Supplementary Table 3. Significant differential splicing events for melanoma cell lines.**

Differential events are defined by q < 0.05, deltaPSI ≥ |0.1|.

**Supplementary Table 4. Significant differential splicing events for ATRT and breast cancer cell lines.** Differential events are defined by q < 0.05, deltaPSI ≥ |0.1|.

**Supplementary Table 5. Summary of significant predicted neopeptides.** Neopeptide candidates were selected based on the product of the -*log*10(%Rank), junction count, and |deltaPSI| value (q < 0.05, deltaPSI ≥ |0.1|, *log*2(TPM) ≥ 3, IJC + SJC ≥ 20, % Rank ≤ 2).

**Supplementary Table 6. Peptides identified from MHCI IP-MS.** Peptides were identified using MSFlagger and MaxQuant (FDR < 0.05).

**Supplementary Table 7. List of antibodies.**

**Supplementary Table 8. List of primers.**

## References

1. Baralle, F. E. & Giudice, J. Alternative splicing as a regulator of development and tissue identity. Nat. Rev. Mol. Cell Biol. 18, 437–451 (2017).

2. Gallego-Paez, L. M. et al. Alternative splicing: the pledge, the turn, and the prestige : The key role of alternative splicing in human biological systems. Hum. Genet. 136, 1015–1042 (2017).

3. Kahles, A. et al. Comprehensive Analysis of Alternative Splicing Across Tumors from 8,705 Patients. Cancer Cell 34, 211–224.e6 (2018).

4. Stanley, R. F. & Abdel-Wahab, O. Dysregulation and therapeutic targeting of RNA splicing in cancer. *Nat*. Cancer 3, 536–546 (2022).

5. Bradley, R. K. & Anczuków, O. RNA splicing dysregulation and the hallmarks of cancer. Nat. Rev. Cancer 23, 135–155 (2023).

6. Ladomery, M. Aberrant alternative splicing is another hallmark of cancer. Int. J. Cell Biol. 2013, 463786 (2013).

7. Oltean, S. & Bates, D. O. Hallmarks of alternative splicing in cancer. Oncogene 33, 5311– 5318 (2014).

8. Desterro, J., Bak-Gordon, P. & Carmo-Fonseca, M. Targeting mRNA processing as an anticancer strategy. Nat. Rev. Drug Discov. 19, 112–129 (2020).

9. Siegfried, Z. & Karni, R. The role of alternative splicing in cancer drug resistance. Curr. Opin. Genet. Dev. 48, 16–21 (2018).

10. Frankiw, L., Baltimore, D. & Li, G. Alternative mRNA splicing in cancer immunotherapy. Nat. Rev. Immunol. 19, 675–687 (2019).

11. Park, J., Park, J. & Chung, Y.-J. Alternative splicing: a new breakthrough for understanding tumorigenesis and potential clinical applications. Genes Genomics 45, 393–400 (2023).

12. Jayasinghe, R. G. et al. Systematic Analysis of Splice-Site-Creating Mutations in Cancer. Cell Rep. 23, 270–281.e3 (2018).

13. Bigot, J. et al. Splicing Patterns in SF3B1-Mutated Uveal Melanoma Generate Shared Immunogenic Tumor-Specific Neoepitopes. Cancer Discov. 11, 1938–1951 (2021).

14. Oka, M. et al. Aberrant splicing isoforms detected by full-length transcriptome sequencing as transcripts of potential neoantigens in non-small cell lung cancer. Genome Biol. 22, 9 (2021).

15. Lu, S. X. et al. Pharmacologic modulation of RNA splicing enhances anti-tumor immunity. Cell 184, 4032–4047.e31 (2021).

16. Matsushima, S. et al. Chemical induction of splice-neoantigens attenuates tumor growth in a preclinical model of colorectal cancer. Sci. Transl. Med. 14, eabn6056 (2022).

17. Xie, N. et al. Neoantigens: promising targets for cancer therapy. Signal Transduct. Target. Ther. 8, 9 (2023).

18. Bhattacharya, S. et al. The methyltransferase SETD2 couples transcription and splicing by engaging mRNA processing factors through its SHI domain. Nat. Commun. 12, 1443 (2021).

19. Duan, L. et al. Histone lysine demethylase KDM4B regulates the alternative splicing of the androgen receptor in response to androgen deprivation. Nucleic Acids Res. 47, 11623– 11636 (2019).

20. Gehring, N. H. & Roignant, J.-Y. Anything but Ordinary – Emerging Splicing Mechanisms in Eukaryotic Gene Regulation. Trends Genet. 37, 355–372 (2021).

21. Inoue, D. et al. Spliceosomal disruption of the non-canonical BAF complex in cancer. Nature 574, 432–436 (2019).

22. Luco, R. F., Allo, M., Schor, I. E., Kornblihtt, A. R. & Misteli, T. Epigenetics in alternative pre-mRNA splicing. Cell 144, 16–26 (2011).

23. Naftelberg, S., Schor, I. E., Ast, G. & Kornblihtt, A. R. Regulation of alternative splicing through coupling with transcription and chromatin structure. Annu. Rev. Biochem. 84, 165– 198 (2015).

24. Shukla, S. et al. CTCF-promoted RNA polymerase II pausing links DNA methylation to splicing. Nature 479, 74–79 (2011).

25. Xiao, R. et al. Pervasive Chromatin-RNA Binding Protein Interactions Enable RNA-Based Regulation of Transcription. Cell 178, 107–121.e18 (2019).

26. Luco, R. F. et al. Regulation of alternative splicing by histone modifications. Science 327, 996–1000 (2010).

27. Luo, C. et al. SRSF2 Regulates Alternative Splicing to Drive Hepatocellular Carcinoma Development. Cancer Res. 77, 1168–1178 (2017).

28. Zhou, X. et al. Splicing factor SRSF1 promotes gliomagenesis via oncogenic splice-switching of MYO1B. J. Clin. Invest. 129, 676–693 (2019).

29. Boddu, P. C. et al. Transcription elongation defects link oncogenic splicing factor mutations to targetable alterations in chromatin landscape. BioRxiv Prepr. Serv. Biol. 2023.02.25.530019 (2023) doi:10.1101/2023.02.25.530019.

30. Rahhal, R. & Seto, E. Emerging roles of histone modifications and HDACs in RNA splicing. Nucleic Acids Res. 47, 4911–4926 (2019).

31. Burbage, M. et al. Epigenetically controlled tumor antigens derived from splice junctions between exons and transposable elements. Sci. Immunol. 8, eabm6360 (2023).

32. Andrés, M. E. et al. CoREST: A functional corepressor required for regulation of neural-specific gene expression. Proc. Natl. Acad. Sci. 96, 9873–9878 (1999).

33. Lee, K., Whedon, S. D., Wang, Z. A. & Cole, P. A. Distinct biochemical properties of the class I histone deacetylase complexes. Curr. Opin. Chem. Biol. 70, 102179 (2022).

34. Wang, Z. A. et al. Diverse nucleosome Site-Selectivity among histone deacetylase complexes. eLife 9, e57663 (2020).

35. Rivera, C. et al. Unveiling RCOR1 as a rheostat at transcriptionally permissive chromatin. Nat. Commun. 13, 1550 (2022).

36. Kalin, J. H. et al. Targeting the CoREST complex with dual histone deacetylase and demethylase inhibitors. Nat. Commun. 9, 53 (2018).

37. Wu, M. et al. The CoREST Repressor Complex Mediates Phenotype Switching and Therapy Resistance in Melanoma. bioRxiv 2020.09.30.320580 (2020) doi:10.1101/2020.09.30.320580.

38. Anastas, J. N. et al. Re-programing Chromatin with a Bifunctional LSD1/HDAC Inhibitor Induces Therapeutic Differentiation in DIPG. Cancer Cell 36, 528–544.e10 (2019).

39. Soukar, I. et al. The CoREST complex is a therapeutic vulnerability in malignant peripheral nerve sheath tumors. BioRxiv Prepr. Serv. Biol. 2024.08.17.607802 (2024) doi:10.1101/2024.08.17.607802.

40. Miller, S. A. et al. Lysine-Specific Demethylase 1 Mediates AKT Activity and Promotes Epithelial-to-Mesenchymal Transition in PIK3CA-Mutant Colorectal Cancer. Mol. Cancer Res. MCR 18, 264–277 (2020).

41. Asmamaw, M. D., He, A., Zhang, L.-R., Liu, H.-M. & Gao, Y. Histone deacetylase complexes: Structure, regulation and function. Biochim. Biophys. Acta Rev. Cancer 1879, 189150 (2024).

42. Garcia-Martinez, L. et al. Endocrine resistance and breast cancer plasticity are controlled by CoREST. Nat. Struct. Mol. Biol. 29, 1122–1135 (2022).

43. Das, S. & Krainer, A. R. Emerging functions of SRSF1, splicing factor and oncoprotein, in RNA metabolism and cancer. Mol. Cancer Res. MCR 12, 1195–1204 (2014).

44. Larsson, C. A., Cote, G. & Quintás-Cardama, A. The changing mutational landscape of acute myeloid leukemia and myelodysplastic syndrome. Mol. Cancer Res. MCR 11, 815–827 (2013).

45. Abramson, J. et al. Accurate structure prediction of biomolecular interactions with AlphaFold 3. Nature 630, 493–500 (2024).

46. Forneris, F., Binda, C., Adamo, A., Battaglioli, E. & Mattevi, A. Structural basis of LSD1-CoREST selectivity in histone H3 recognition. J. Biol. Chem. 282, 20070–20074 (2007).

47. Kim, S.-A., Zhu, J., Yennawar, N., Eek, P. & Tan, S. Crystal Structure of the LSD1/CoREST Histone Demethylase Bound to Its Nucleosome Substrate. Mol. Cell 78, 903–914.e4 (2020).

48. Maji, D. et al. Representative cancer-associated U2AF2 mutations alter RNA interactions and splicing. J. Biol. Chem. 295, 17148–17157 (2020).

49. Glasser, E. et al. Pre-mRNA splicing factor U2AF2 recognizes distinct conformations of nucleotide variants at the center of the pre-mRNA splice site signal. Nucleic Acids Res. 50, 5299–5312 (2022).

50. Bonnal, S. C., López-Oreja, I. & Valcárcel, J. Roles and mechanisms of alternative splicing in cancer — implications for care. Nat. Rev. Clin. Oncol. 17, 457–474 (2020).

51. Arozarena, I. & Wellbrock, C. Phenotype plasticity as enabler of melanoma progression and therapy resistance. Nat. Rev. Cancer 19, 377–391 (2019).

52. Wang, Y. et al. rMATS-turbo: an efficient and flexible computational tool for alternative splicing analysis of large-scale RNA-seq data. Nat. Protoc. 19, 1083–1104 (2024).

53. Li, Y. I. et al. Annotation-free quantification of RNA splicing using LeafCutter. Nat. Genet. 50, 151–158 (2018).

54. Li, G. et al. Splicing neoantigen discovery with SNAF reveals shared targets for cancer immunotherapy. Sci. Transl. Med. 16, eade2886 (2024).

55. Rybak, J.-N., Roesli, C., Kaspar, M., Villa, A. & Neri, D. The extra-domain A of fibronectin is a vascular marker of solid tumors and metastases. Cancer Res. 67, 10948–10957 (2007).

56. Kim, Y.-E. et al. RBM47-regulated alternative splicing of TJP1 promotes actin stress fiber assembly during epithelial-to-mesenchymal transition. Oncogene 38, 6521–6536 (2019).

57. Zhang, Y. et al. OncoSplicing: an updated database for clinically relevant alternative splicing in 33 human cancers. Nucleic Acids Res. 50, D1340–D1347 (2022).

58. Flemington, E. K. et al. SpliceTools, a suite of downstream RNA splicing analysis tools to investigate mechanisms and impact of alternative splicing. Nucleic Acids Res. 51, e42–e42 (2023).

59. Reynisson, B., Alvarez, B., Paul, S., Peters, B. & Nielsen, M. NetMHCpan-4.1 and NetMHCIIpan-4.0: improved predictions of MHC antigen presentation by concurrent motif deconvolution and integration of MS MHC eluted ligand data. Nucleic Acids Res. 48, W449– W454 (2020).

60. Sarkizova, S. et al. A large peptidome dataset improves HLA class I epitope prediction across most of the human population. Nat. Biotechnol. 38, 199–209 (2020).

61. Li, G., Iyer, B., Prasath, V. B. S., Ni, Y. & Salomonis, N. DeepImmuno: deep learning-empowered prediction and generation of immunogenic peptides for T-cell immunity. Brief. Bioinform. 22, bbab160 (2021).

62. Aran, D. et al. Reference-based analysis of lung single-cell sequencing reveals a transitional profibrotic macrophage. Nat. Immunol. 20, 163–172 (2019).

63. El Marabti, E. & Younis, I. The Cancer Spliceome: Reprograming of Alternative Splicing in Cancer. Front. Mol. Biosci. 5, 80 (2018).

64. Floro, J. et al. SDE2 is an essential gene required for ribosome biogenesis and the regulation of alternative splicing. Nucleic Acids Res. 49, 9424–9443 (2021).

65. Shen, S.-M. et al. Nuclear PTEN safeguards pre-mRNA splicing to link Golgi apparatus for its tumor suppressive role. Nat. Commun. 9, 2392 (2018).

66. Glasser, E., Agrawal, A. A., Jenkins, J. L. & Kielkopf, C. L. Cancer-Associated Mutations Mapped on High-Resolution Structures of the U2AF2 RNA Recognition Motifs. Biochemistry 56, 4757–4761 (2017).

67. Kang, H.-S. et al. An autoinhibitory intramolecular interaction proof-reads RNA recognition by the essential splicing factor U2AF2. Proc. Natl. Acad. Sci. U. S. A. 117, 7140–7149 (2020).

68. Schneider-Poetsch, T., Chhipi-Shrestha, J. K. & Yoshida, M. Splicing modulators: on the way from nature to clinic. J. Antibiot. (Tokyo) 74, 603–616 (2021).

69. Feustel, K. & Falchook, G. S. Protein Arginine Methyltransferase 5 (PRMT5) Inhibitors in Oncology Clinical Trials: A review. J. Immunother. Precis. Oncol. 5, 58–67 (2022).

70. Hwang, J. W., Cho, Y., Bae, G.-U., Kim, S.-N. & Kim, Y. K. Protein arginine methyltransferases: promising targets for cancer therapy. Exp. Mol. Med. 53, 788–808 (2021).

71. Westcott, P. M. K. et al. Low neoantigen expression and poor T-cell priming underlie early immune escape in colorectal cancer. *Nat*. Cancer 2, 1071–1085 (2021).

72. Samstein, R. M. et al. Tumor mutational load predicts survival after immunotherapy across multiple cancer types. Nat. Genet. 51, 202–206 (2019).

73. Topalian, S. L. et al. Neoadjuvant immune checkpoint blockade: A window of opportunity to advance cancer immunotherapy. Cancer Cell 41, 1551–1566 (2023).

74. Palmeri, M. et al. Real-world application of tumor mutational burden-high (TMB-high) and microsatellite instability (MSI) confirms their utility as immunotherapy biomarkers. ESMO Open 7, 100336 (2022).

75. Truong, A. S. et al. Entinostat induces antitumor immune responses through immune editing of tumor neoantigens. J. Clin. Invest. 131, e138560 (2021).

76. van Bergen, M. G. J. M. & van der Reijden, B. A. Targeting the GFI1/1B-CoREST Complex in Acute Myeloid Leukemia. Front. Oncol. 9, 1027 (2019).

77. Feldman, J., Goldwasser, R., Mark, S., Schwartz, J. & Orion, I. A Mathematical Model for Tumor Volume Evaluation using Two-Dimensions. J. Appl. Quanitative Methods 4, (2009).

78. Aguilan, J. T., Kulej, K. & Sidoli, S. Guide for protein fold change and *p* -value calculation for non-experts in proteomics. *Mol*. Omics 16, 573–582 (2020).

79. Lee, K. et al. Uncoupling histone modification crosstalk by engineering lysine demethylase LSD1. Nat. Chem. Biol. (2024) doi:10.1038/s41589-024-01671-9.

80. Punjani, A., Rubinstein, J. L., Fleet, D. J. & Brubaker, M. A. cryoSPARC: algorithms for rapid unsupervised cryo-EM structure determination. Nat. Methods 14, 290–296 (2017).

81. Punjani, A., Zhang, H. & Fleet, D. J. Non-uniform refinement: adaptive regularization improves single-particle cryo-EM reconstruction. Nat. Methods 17, 1214–1221 (2020).

82. Pettersen, E. F. et al. UCSF ChimeraX: Structure visualization for researchers, educators, and developers. Protein Sci. Publ. Protein Soc. 30, 70–82 (2021).

83. Emsley, P., Lohkamp, B., Scott, W. G. & Cowtan, K. Features and development of Coot. Acta Crystallogr. D Biol. Crystallogr. 66, 486–501 (2010).

84. Liebschner, D. et al. Macromolecular structure determination using X-rays, neutrons and electrons: recent developments in Phenix. Acta Crystallogr. Sect. Struct. Biol. 75, 861–877 (2019).

85. Williams, C. J. et al. MolProbity: More and better reference data for improved all-atom structure validation. Protein Sci. Publ. Protein Soc. 27, 293–315 (2018).

86. Bolger, A. M., Lohse, M. & Usadel, B. Trimmomatic: a flexible trimmer for Illumina sequence data. Bioinformatics 30, 2114–2120 (2014).

87. Dobin, A. et al. STAR: ultrafast universal RNA-seq aligner. Bioinformatics 29, 15–21 (2013).

88. Liao, Y., Smyth, G. K. & Shi, W. The Subread aligner: fast, accurate and scalable read mapping by seed-and-vote. Nucleic Acids Res. 41, e108–e108 (2013).

89. Love, M. I., Huber, W. & Anders, S. Moderated estimation of fold change and dispersion for RNA-seq data with DESeq2. Genome Biol. 15, 550 (2014).

90. Subramanian, A. et al. Gene set enrichment analysis: A knowledge-based approach for interpreting genome-wide expression profiles. Proc. Natl. Acad. Sci. 102, 15545–15550 (2005).

91. Kanehisa, M., Furumichi, M., Tanabe, M., Sato, Y. & Morishima, K. KEGG: new perspectives on genomes, pathways, diseases and drugs. Nucleic Acids Res. 45, D353– D361 (2017).

92. Liberzon, A. et al. The Molecular Signatures Database Hallmark Gene Set Collection. Cell Syst. 1, 417–425 (2015).

93. Gao, J. et al. Integrative analysis of complex cancer genomics and clinical profiles using the cBioPortal. Sci. Signal. 6, pl1 (2013).

94. Ficarro, S. B. et al. Improved electrospray ionization efficiency compensates for diminished chromatographic resolution and enables proteomics analysis of tyrosine signaling in embryonic stem cells. Anal. Chem. 81, 3440–3447 (2009).

95. Kong, A. T., Leprevost, F. V., Avtonomov, D. M., Mellacheruvu, D. & Nesvizhskii, A. I. MSFragger: ultrafast and comprehensive peptide identification in mass spectrometry-based proteomics. Nat. Methods 14, 513–520 (2017).

96. Tyanova, S., Temu, T. & Cox, J. The MaxQuant computational platform for mass spectrometry-based shotgun proteomics. Nat. Protoc. 11, 2301–2319 (2016).

97. Hao, Y. et al. Dictionary learning for integrative, multimodal and scalable single-cell analysis. Nat. Biotechnol. 42, 293–304 (2024).

98. McGinnis, C. S., Murrow, L. M. & Gartner, Z. J. DoubletFinder: Doublet Detection in Single-Cell RNA Sequencing Data Using Artificial Nearest Neighbors. Cell Syst. 8, 329–337.e4 (2019).

99. Hafemeister, C. & Satija, R. Normalization and variance stabilization of single-cell RNA-seq data using regularized negative binomial regression. Genome Biol. 20, 296 (2019).

100. Zappia, L. & Oshlack, A. Clustering trees: a visualization for evaluating clusterings at multiple resolutions. GigaScience 7, giy083 (2018).

101. Finak, G. et al. MAST: a flexible statistical framework for assessing transcriptional changes and characterizing heterogeneity in single-cell RNA sequencing data. Genome Biol. 16, 278 (2015).

102. Korotkevich, G. et al. Fast gene set enrichment analysis. Preprint at 10.1101/060012 (2021).

103. Kuznetsova, A., Brockhoff, P. B. & Christensen, R. H. B. lmerTest Package: Tests in Linear Mixed Effects Models. J. Stat. Softw. 82, (2017).

104. Hwang, J. Y. et al. rMAPS2: an update of the RNA map analysis and plotting server for alternative splicing regulation. Nucleic Acids Res. 48, W300–W306 (2020).

