## Supplementary figures and images for "CoREST Complex Inhibition Alters RNA Splicing to Promote Neoantigen Expression and Enhance Tumor Immunity"

### Supplemental Figures

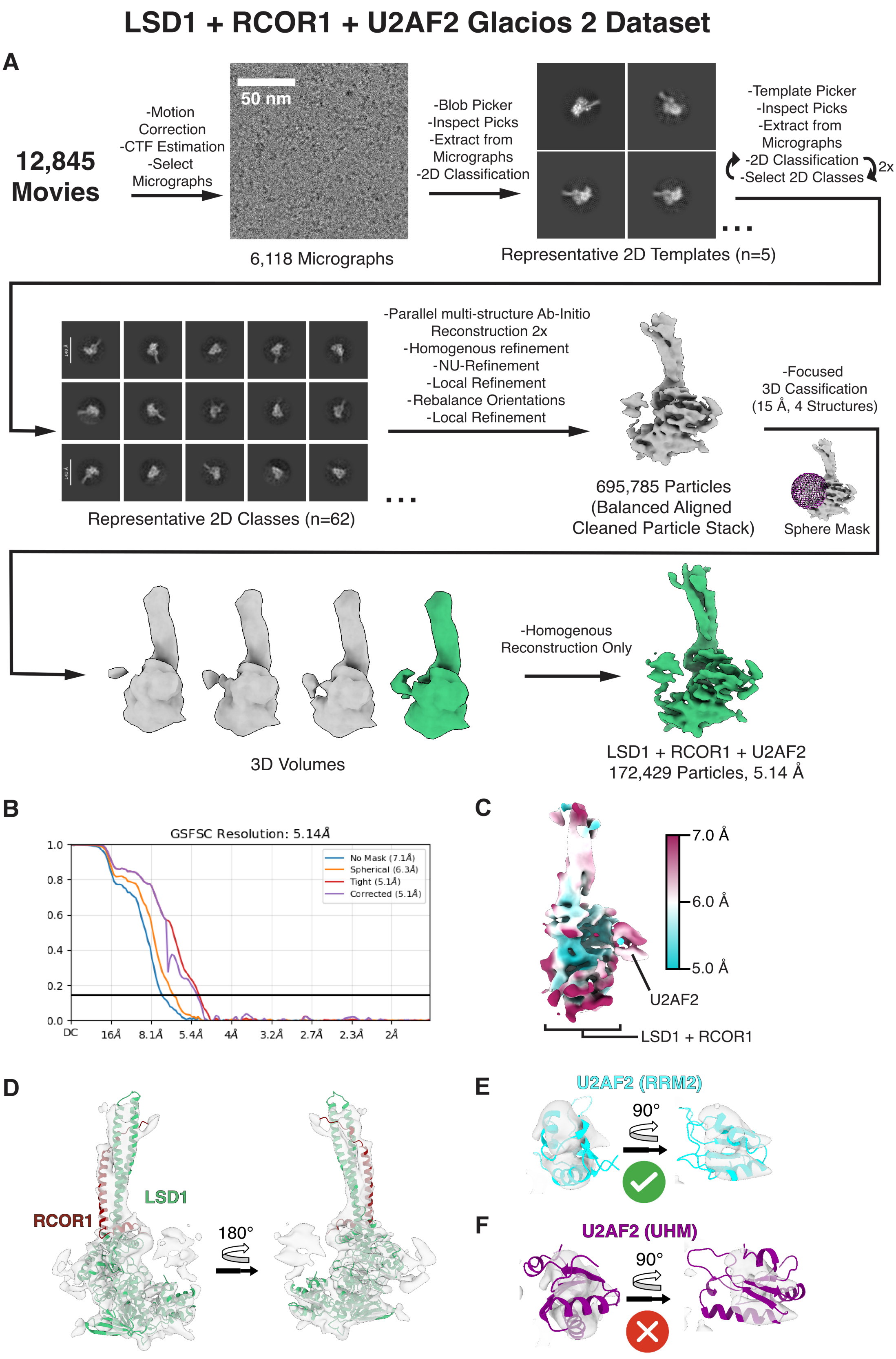

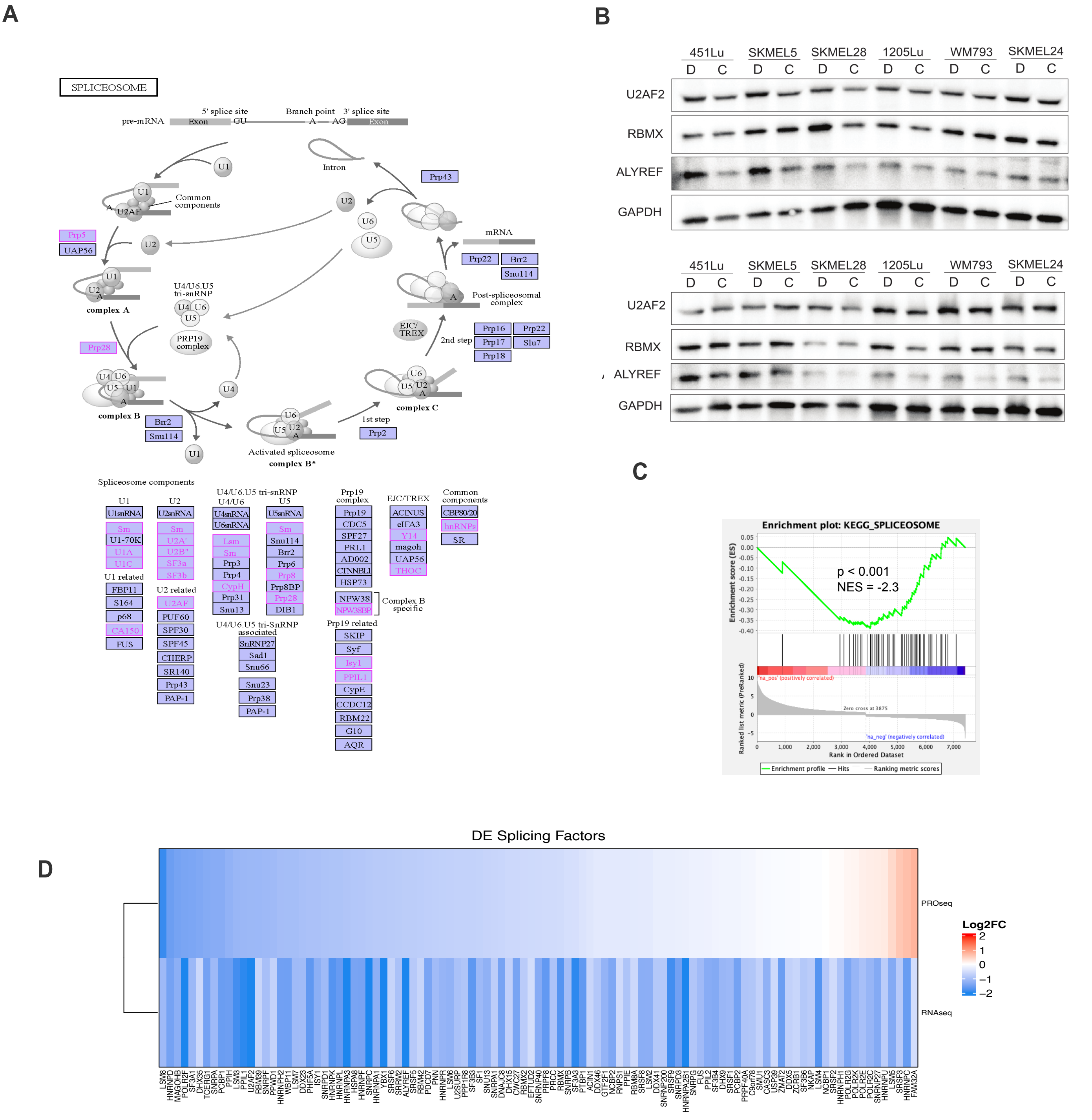

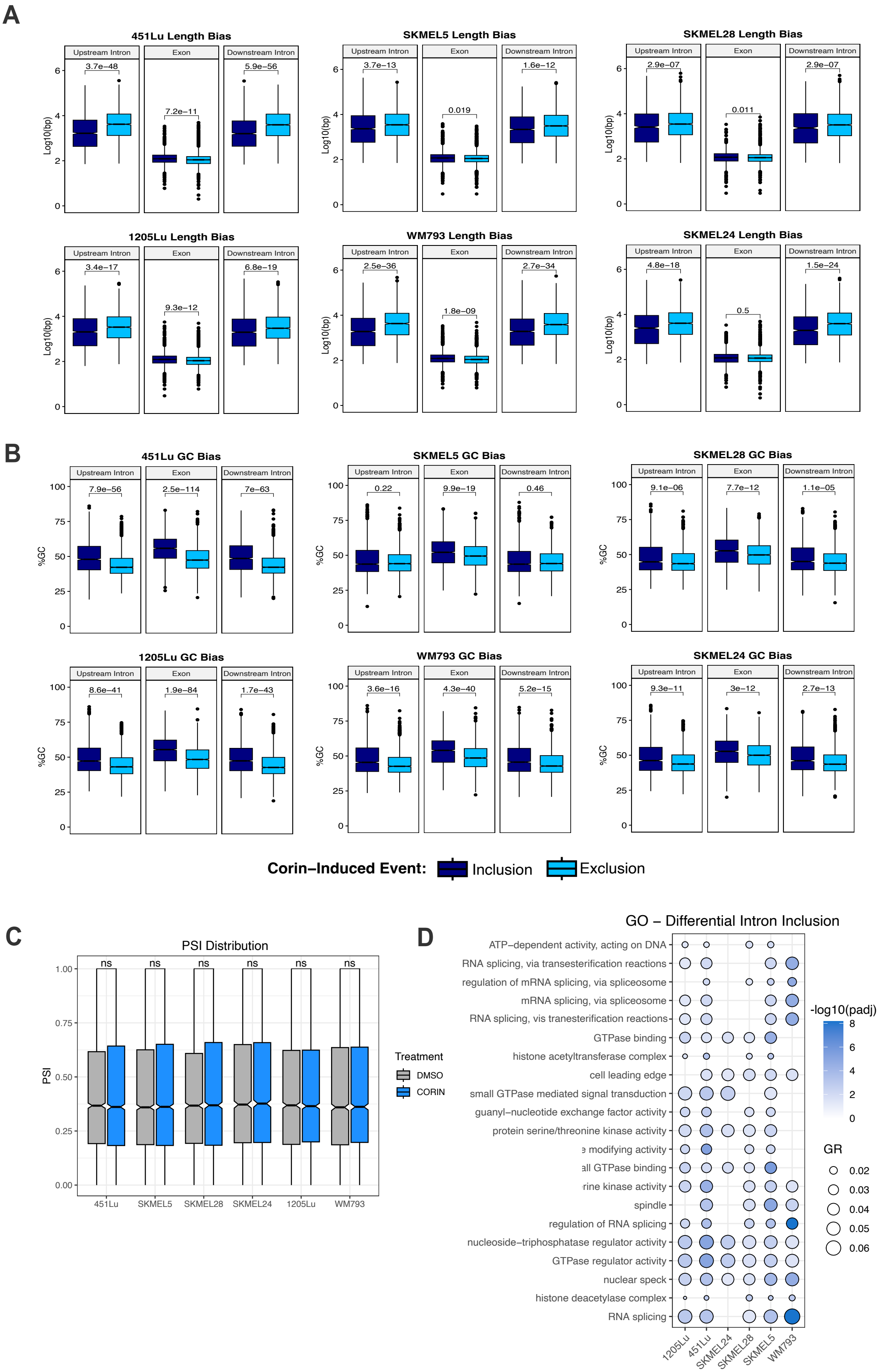

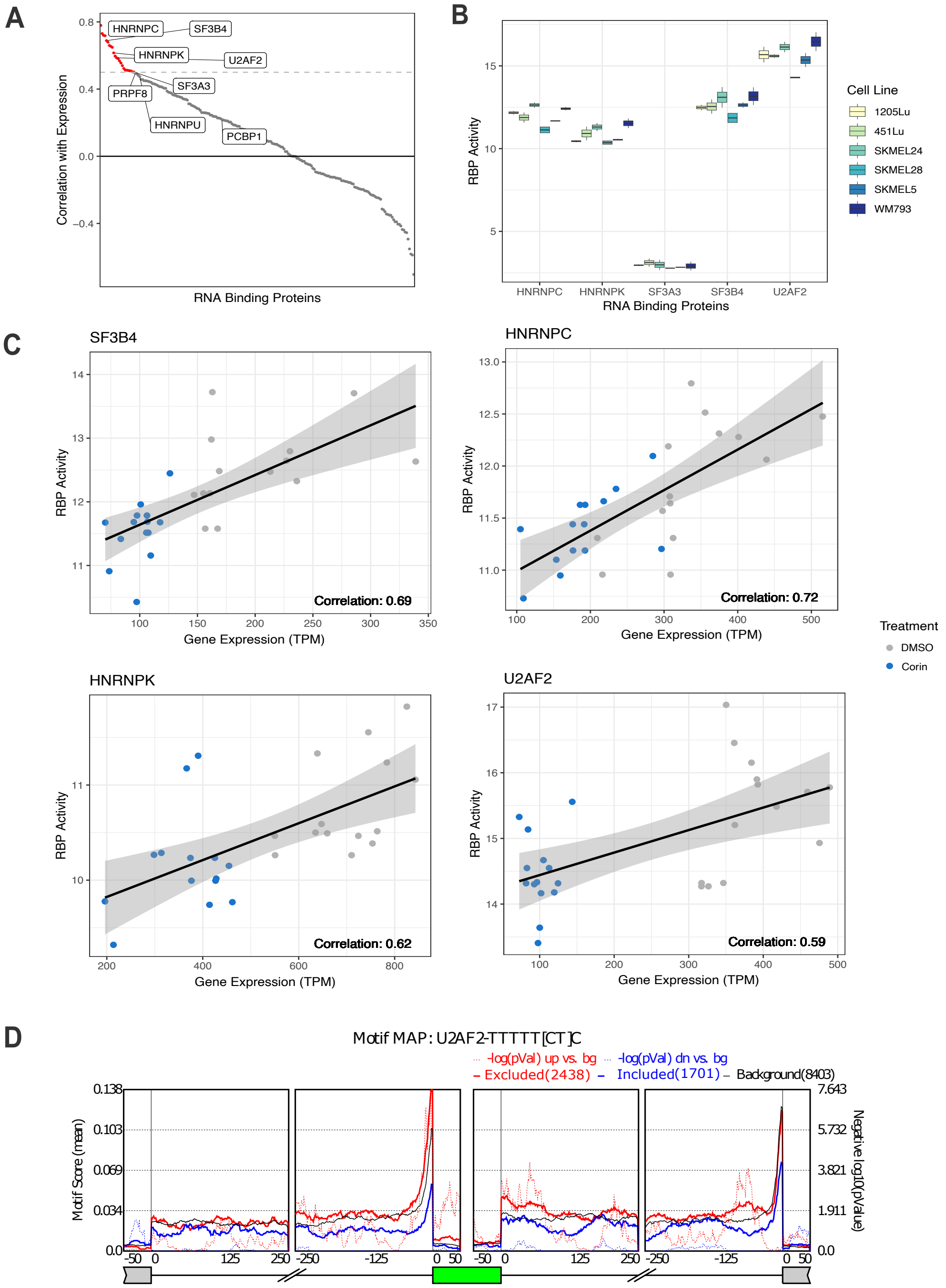

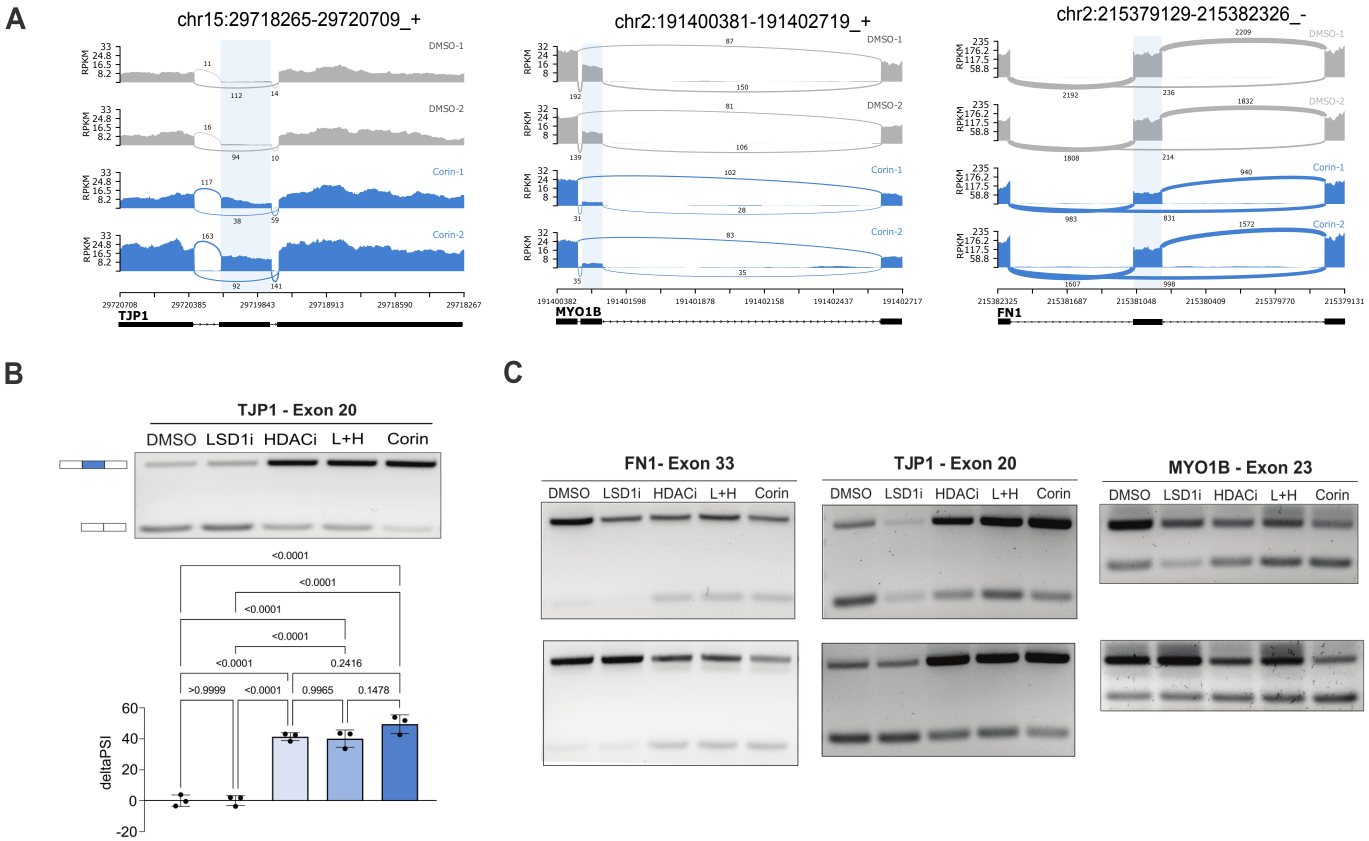

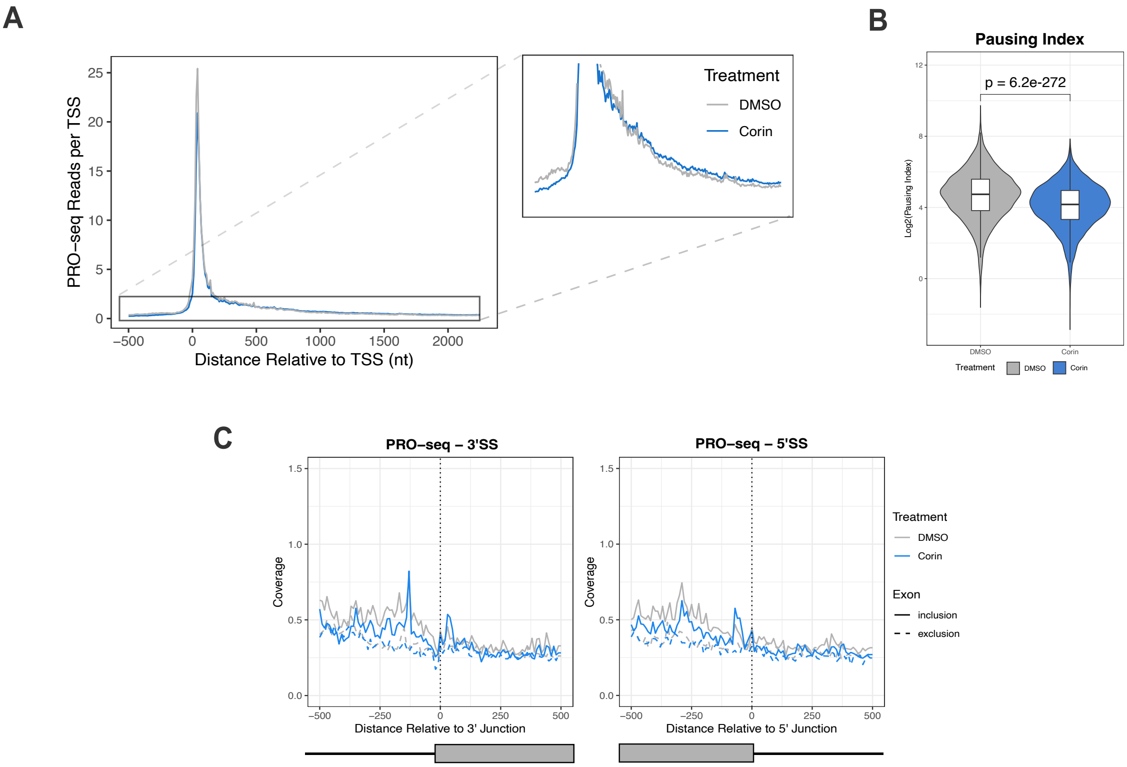

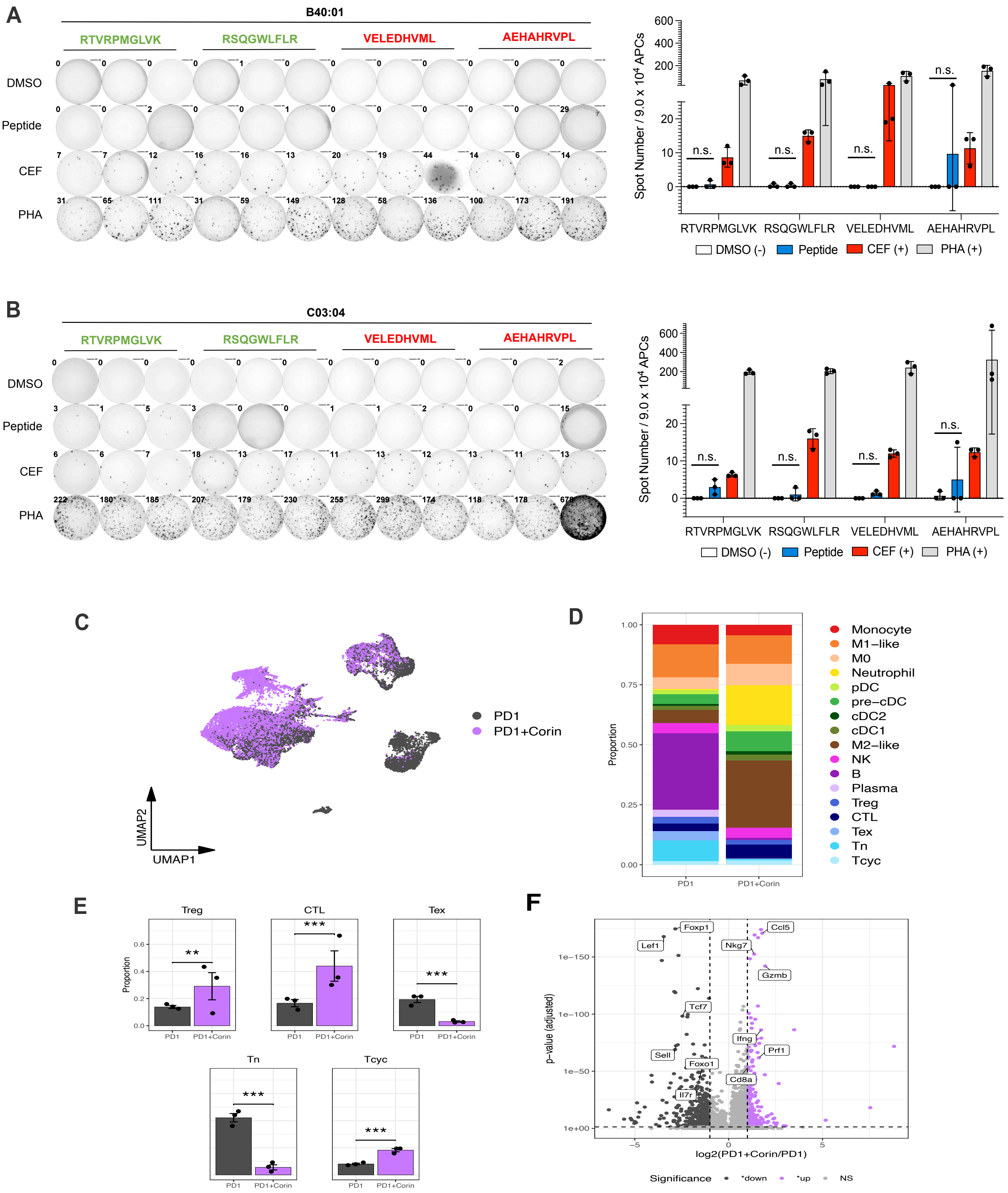
